# Mesopelagic fish responses to Pleistocene climatic variability in the Eastern Mediterranean and implications for the biological pump

**DOI:** 10.1101/2024.12.28.630586

**Authors:** Konstantina Agiadi, Iuliana Vasiliev, Antoine Vite, Stergios Zarkogiannis, Alba Fuster-Alonso, Jorge Mestre-Tomás, Efterpi Koskeridou, Frédéric Quillévéré

## Abstract

Mesopelagic fishes play a crucial role in the global carbon cycle through their diel vertical migration (DVM), but the impacts of neither natural nor anthropogenic climate change on DVM patterns are currently known. Studying the geological past can elucidate changes in DVM patterns under swelling climate pressure and allow estimating the impacts of the current climate crisis. We present a multi-proxy, ecosystem-level assessment of paleoenvironmental changes in the Eastern Mediterranean during the Middle Pleistocene (marine isotope stages MIS 23–18; 923–756 ka B.P.) and use the carbon and oxygen isotopic composition of fossil fish otoliths to assess the impacts of these changes on DVM and their possible implications on the biological pump. Temperature was the primary driver of ecosystem change during MIS 21 interglacial, whereas productivity became a dominant factor in MIS 19 interglacial. Responses of organisms throughout the water column varied. Our results indicate increased productivity across trophic levels during MIS 19, which affected foraminiferal biomasses, but did not inhibit fish DVM. In contrast, the early MIS 21 warming led to a reduction in DVM by mesopelagic fishes and consequently a drop in biological pump efficiency.

## Introduction

In pelagic ecosystems, mesopelagic fishes form the link between trophic levels and play an essential role in the biological carbon pump due to their diel vertical migration (DVM). DVM is the daily movement of several marine organisms, including mesopelagic fishes of the family Myctophidae (commonly known as lanternfishes), between the twilight (day-time) and the euphotic (night-time) zones (Brierley, 2014). Because these fishes feed at the surface and excrete and respire carbon at great depths in the mesopelagic zone of the oceans, their DVM leads to a net drawdown of carbon from the surface waters. According to some estimates, DVM contributes approximately 20% of the biological carbon pump in modern oceans, with more than 1 PgC/y being actively transported below the euphotic zone by all migrating organisms (Aumont et al., 2018). Compared with any other process of the biological carbon pump, carbon is directly injected deeper through the DVM (Aumont et al., 2018). As a result, the contribution of DVM to carbon sequestration accounts for more than 50% of the global total, and mesopelagic fishes contribute half of this total (Aksnes et al., 2023; Pinti et al., 2023). The capacity of myctophids to perform DVM varies by species and is affected by seawater conditions, as well as food availability and type (Olivar et al., 2018).

Nothing is known about DVM in the geological past, despite its likely significant contribution to carbon sequestration and paleoclimate variability. Fish, in particular, respond swiftly to unfavorable environmental conditions by adjusting their geographic and depth distribution ranges (Perry et al., 2005), but changes in temperature, salinity, oxygenation and the availability of food over time lead to eco-evolutionary shifts in the metabolic rates of these fishes, as well as in their size at maturity and life-history patterns (Gallo and Levin, 2016; Lindmark et al., 2022; Rountrey et al., 2014).

We investigate here changes in the DVM patterns of two of the most common mesopelagic fishes worldwide, *Hygophum benoiti* and *Ceratoscopelus maderensis*, between 923 and 756 ka B.P. (marine isotope stages MIS 23–MIS 18), which was an interval characterized by strong paleoclimatic changes (Elderfield et al., 2012; Lisiecki and Raymo, 2005b). We obtain the thermohaline gradient, the fish depth distribution and their DVM patterns in the Eastern Mediterranean based on the oxygen isotopic compositions of foraminifera shells and fish otoliths. In addition, we infer changes in fish metabolism based on the carbon stable isotopic composition of the same material. By reconstructing sea surface temperature (SST), salinity (SSS), primary and secondary production (foraminiferal biomass, ostracod and sponge spicule counts), we are able to investigate the relationship between paleoclimatic and paleoceanographic changes, and the mesopelagic fish DVM patterns.

### The Pleistocene Mediterranean climate and marine ecosystem

The Pleistocene Mediterranean has been the topic of numerous studies because it was at the time a hotspot of ancient human population change associated with natural climate changes (Margari et al., 2023). For the Early and Middle Pleistocene, SST and primary productivity changes have been reconstructed with high temporal resolution in the Western and Central Mediterranean (Marino et al., 2020, 2022; Martínez-Dios et al., 2021; Nomade et al., 2019; Quivelli et al., 2021) and the southwestern Iberian margin (Girone et al., 2023; Rodrigues et al., 2017; Sánchez Goñi et al., 2023), providing up to millennial-scale paleoclimate reconstructions with even records of short-term strong cooling events attributed to the reduction of the Atlantic Meridional Overturning Circulation. However, very few studies have been conducted in the Eastern Mediterranean (Eichner et al., 2024a; Incarbona et al., 2022; Quillévéré et al., 2016, 2019). The Eastern Mediterranean is currently affected by particularly much drier and ∼4 °C warmer climate than the Western Mediterranean (Minnett et al., 2019). In addition, the riverine influx only partly counterbalances, the negative water budget of the region promoted by intense evaporation, and therefore much saltier sea surface conditions (> 39) prevail there than in the Western Mediterranean (∼36.5).

Climate change affects the size and metabolism of marine organisms (Daufresne et al., 2009; Ohlberger, 2013), and consequently, marine ecosystem structure and function (Hildrew et al., 2007). Although some aspects of the response of marine organisms to the paleoclimatic changes that occurred in the Mediterranean during the Pleistocene have been studied so far, such as the composition of the calcareous nannoplankton (Marino et al., 2008, 2020), planktonic foraminifera (Quillévéré et al., 2019; Quivelli et al., 2021), and fish communities (Agiadi et al., 2011, 2018; Girone and Varola, 2001), as well as the body size of bivalves (Porz et al., 2024) and fishes (Agiadi et al., 2023), prevailing questions remain about secondary production by zooplankton and zoobenthos, and the metabolic response of higher-trophic–level organisms.

## Methods

### Study area and sampling

The material used in this study originates from the Lindos Bay Formation exposed at Lardos, northeastern Rhodes, Greece (Fig. 1). On the island of Rhodes, Mesozoic–Paleogene limestones are overlain by marine siliciclastic–carbonate Pleistocene deposits. Since the Pliocene, the compressive regime of this area of the Hellenic forearc and the concomitant opening of the > 4000-m–deep Rhodes Basin have triggered rapid vertical tectonic motions, resulting in the deposition and uplift of Pleistocene marine sediments that are now exposed along the eastern coast of the island (Cornée et al., 2006, 2019; Van Hinsbergen et al., 2007). The Lower–Middle Pleistocene hemipelagic sediments of the Lindos Bay Formation sampled for this study were deposited at bathyal paleodepths during the maximum flooding of this tectonically driven transgression–regression cycle (Cornée et al., 2019; Milker et al., 2017; Quillévéré et al., 2019). This occurrence of Pleistocene hemipelagic sediments accessible on land is unique for the Eastern Mediterranean, providing a unique opportunity for studying the Early and Middle Pleistocene marine faunas of the region (Quillévéré et al., 2016).

**Figure 1.**
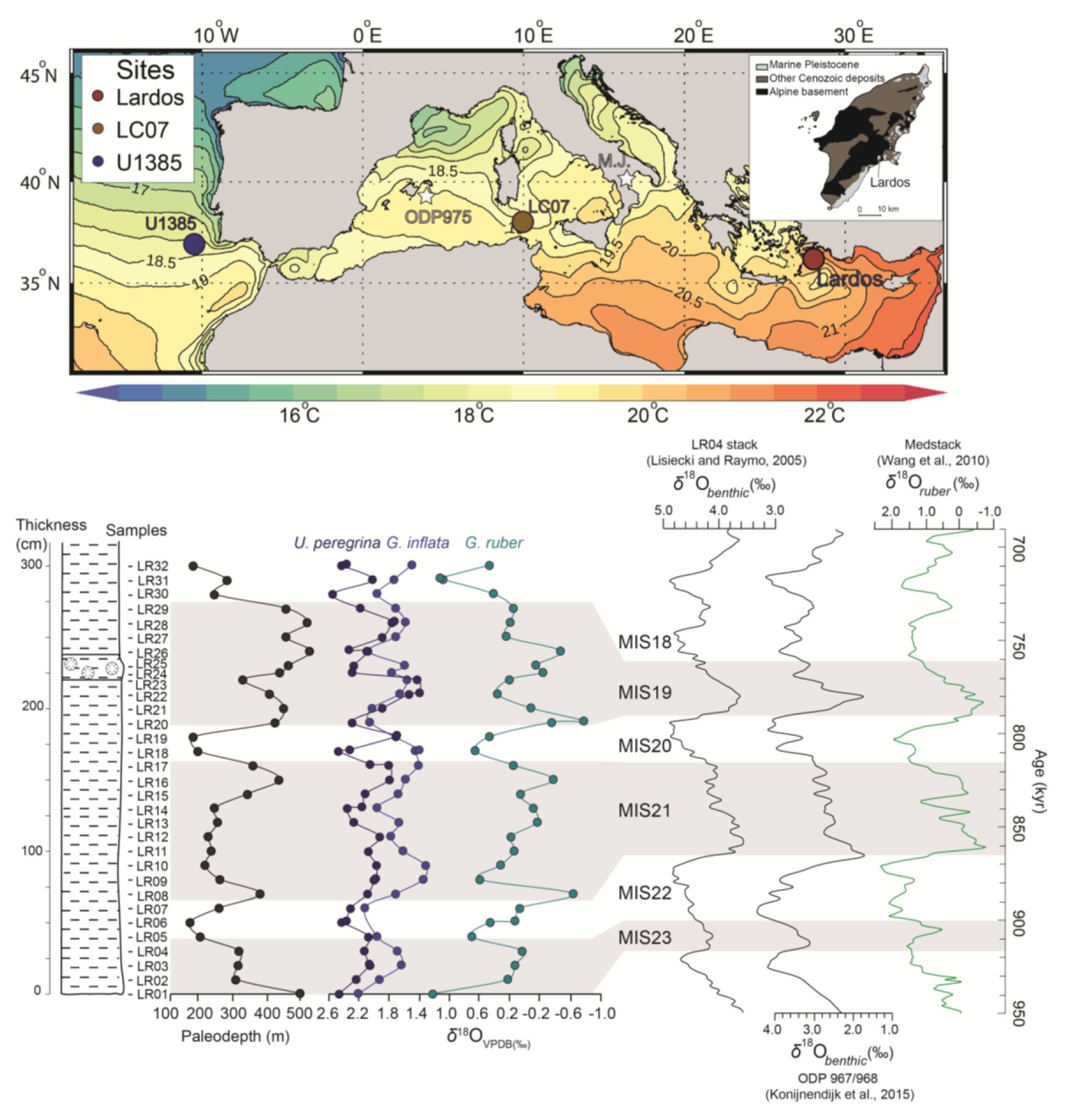
Map of the Mediterranean region with present-day SST (Minnett et al., 2019) and the location of Lardos (our study area), LC07 (Martínez-Dios et al., 2021), U1385 (Rodrigues et al., 2017), ODP975 (Quivelli et al., 2020, 2021), and M.J. Montalbano Jonico (Marino et al., 2020; Peral et al., 2020). Paleodepth estimates based on the *%P/(B+P)* ratio and proposed correlation between foraminifera *δ*^18^O records of the Lardos section and open ocean *δ*^18^O reference records: the LR04 benthic global *δ*^18^O stack (Lisiecki and Raymo, 2005), the benthic Eastern Mediterranean *δ*^18^O compilation data from Ocean Drilling Program (ODP) Sites 967 and 968 (Konijnendijk et al., 2015), and the planktonic (*Globigerinoides ruber*) *δ*^18^O Medstack (Wang et al., 2010). The paleodepth changes (with a glacio-eustatic component) and the coincidence of successive positive and negative excursions in the *δ*^18^O records allow refining the chronostratigraphy previously published for the section (Titschack et al., 2013). This revised age model reveals the identification of six distinct isotopic stages between MIS 23 and MIS 18, including three interglacials (gray areas) and three glacials (white areas).

The studied section (36°05’19.5”N, 28°00’35.9”E) is located at the northern flank of a hill positioned ∼900 m southwest of the village of Lardos. The total section comprises 9.3 meters of marine marls and silts, which belong to the Lindos Bay Formation followed by the lowermost part of the Arkhangellos Formation (Titschack et al. 2013). In this work, we targeted the 2.4 m lowest marls of the section corresponding to the Lindos Bay Formation, which had been previously studied by Titschack et al. (2013). Additionally, we dug a pit beneath them, extending the section by 0.7 m. Thirty-two samples were obtained every 10 cm, after the outcrop surface was refreshed, from the newly exposed 3.1 m.

### Chronostratigraphic background

We revised the age model for Lardos of Titschack et al. (2013), based on our new analyses and considering the model of Eichner et al. (2024b), as explained below:

A chronostratigraphic framework of Lardos section was established by Titschack et al. (2013) on the basis of nannoplankton and foraminiferal biostratigraphic analyses, oxygen isotopic compositions of surface-dwelling planktonic foraminifera, and the U^234^/U^238^ dating of deep-water coral fragments. The proposed age model suggested that deposition of the entire Lardos section occurred between ∼860 and ∼330 ka B.P. Eichner et al. (2024b) recently proposed another age model, in which deposition took place between 751 and 427 ka B.P. Here, we favor the age model of Titschack et al. (2013), which has been applied to several studies of the section (Agiadi et al., 2018, 2023; Porz et al., 2024). Titschack et al. (2013) analyzed three *Desmophyllum pertusum* coral fragments collected from two debris flows cropping out in the lower part of the section. The two fragments from the∼25 cm-thick lowermost debris flow at ∼1.6 m section height of the original section (now at 2.3 m height; Fig. 1) yielded mean ages of 755.2 ± 15.5 ka and 756.0 ± 17.5 ka. A third fragment from the uppermost debris flow level (located ∼2 meters above the part of the section we analyzed in this study) yielded a mean age of 688.9 ± 15.5 ka. Both age models (Eichner et al., 2024b; Titschack et al., 2013) are supported by the same calcareous nannofossil biostratigraphic data and the continuous occurrence of benthic foraminifera *Stilostomella* sp. in the lower part of the section. The model of Titschack et al. (2013) was validated further by Agiadi et al. (2023), who provided a new ^234^U/^238^U dating for an additional *D. pertusum* fragment collected from the lowermost debris flow horizon, yielding a mean age of 744 ka (95% confidence intervals). In contrast, the age model proposed by Eichner et al. (2024b) entirely relies on peak-to-peak visual correlations between their benthic foraminiferal *δ*^18^O data and the global ocean LR04 record (Lisiecki and Raymo, 2005a), but disregarded the coral ages produced by Titschack et al. (2013) as tie points (the fourth ^234^U/^238^U age published by Agiadi et al. (2023) was not cited at all). Eichner et al. (2024b) hypothesized that the coral fragments analyzed by Titschack et al. (2013) from two different debris flow levels, had been systematically reworked or that the ^234^U/^238^U dating did not meet the necessary criteria and an open system could have biased the method (producing older ages). However, there is no tangible evidence of reworking of the coral fragments collected from different levels in such type of upper bathyal environments and hemipelagic deposits. In addition, it is highly unlikely that all four coral fragments collected from two different levels yielded nearly constant age offsets of ∼50–60 ka. Consequently, we consider that the age model of Titschack et al. (2013) was indeed closer to reality.

The local oxygen isotope stratigraphy and paleodepth estimates produced in this study were used to refine the chronostratigraphy of the studied deposits at Lardos that was published by Titschack et al. (2013). In addition, by comparing new local *δ*^18^Ο compositions of monospecific samples of benthic and planktonic foraminifera to those occurring in the global ocean (Lisiecki and Raymo, 2005a) and Mediterranean reference records (Konijnendijk et al., 2015; Wang et al., 2010), we refined the location, in the section, of climatic cycle terminations within the studied interval.

### Paleodepth estimates

The 32 sediment samples were dried in an oven, weighed and placed in water and hydrogen peroxide for five hours on a shaker table. The samples were subsequently sieved through a 63-μm mesh sieve and dried in an oven at 60 °C, after which the residues were weighed. The depositional depth (paleodepth) through the 3.1 m sedimentary unit was assessed by counting in each sample the number of planktonic foraminifera *P* relative to the total number of foraminifera *(P+B)* (up to 300 specimens) in representative splits of the >125 µm size fraction. Indeed, the planktonic/benthic ratio *%P(P+B)* typically increases with paleodepth or distance from shorelines (Gibson, 1989). We estimated the paleodepth *D* via the transfer function developed by van der Zwaan et al. (1990) (Eq. 1):

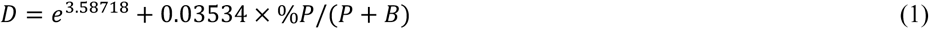

Several factors, such as food availability and oxygen content of seawater (Jorissen et al., 1995), control the depth distributions of benthic foraminifera potentially affecting paleodepth estimates. We consequently followed the methodology proposed by van der Zwaan et al. (1990), discarding from the counts the infaunal benthic genera *Uvigerina*, *Bolivina*, *Bulimina*, *Globobulimina* and *Fursenkoina*. These taxa favor organic-rich and oxygen-deficient environments and their abundances depend mostly on food availability rather than depth.

### Organic matter content and sources

Productivity proxies produce different results depending on the setting and origin of the organic matter (Rühlemann et al., 1996). Relative changes in total productivity were estimated from the TOC content and the bulk *δ*^13^C, considering the sedimentary C/N ratio to distinguish between marine and terrestrial organic matter (El Frihmat et al., 2015) (full measurements available as the dataset Agiadi et al. (2024b). In addition, we have to take into account the sedimentation rate, that may differ between glacials and interglacials. Sample preparation included grinding, removal of inorganic carbon (using 10 % HCl for 24 h at 40 °C), centrifugation (4 × at 2800 to 3000 rpm for 4 to 8 min), and sample drying (24 h at 40 °C). About 8 to 10 mg of sample were analyzed with a Flash EA 1112 that was coupled to a Thermo Fisher MAT 253 gas-source isotope ratio mass spectrometer at the Goethe University–SBiK-F Joint Stable Isotope Facility. An uncertainty of 0.2 ‰ was indicated for measured *δ*^13^C_TOC_ values by analyzing USGS 24 and IAEA-CH-7 standard materials on a daily basis and replicate measurements of reference materials and samples. The total organic carbon and total nitrogen concentrations (in %) have been calculated by relating the signal size of the samples and the averaged signal size of the daily standards (USGS 24, n = 8). A typical error of ∼0.5 % based on mass spectrometric analysis and maximum differences of ∼7 % in TOC contents of replicate measurements (including weighing uncertainties) was detected.

To evaluate the relative input of terrestrial versus marine organic matter we used Branched Isoprenoid Tetraether (BIT) index (Hopmans et al., 2004; Weijers et al., 2006), a biomarker-based proxy calculated as (Eq. 2):

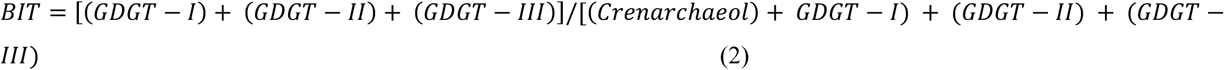

where *GDGT-I*, *GDGT-II* and *GDGT-III* are branched (br)GDGTs, which are components of bacterial cell membranes (particularly from Acidobacteria found in terrestrial, estuarine, and marine environments (Sinninghe Damsté et al., 2011; Weijers et al., 2006). *Crenarchaeol* is an isoGDGT and, together with brGDGTs, was analyzed and isolated as explained in section 2.4. The BIT data are provided in the dataset Agiadi et al. (2024b).

### Oxygen and carbon isotopic analyses

Stable oxygen (*δ*^18^Ο) and carbon (*δ*^13^C) isotopic compositions were first measured on the shells of benthic (*Uvigerina peregrina*) and planktonic foraminifera (*Globigerinoides ruber, Globoconella inflata*), and fish otoliths (*Hygophum benoiti*, *Ceratoscopelus maderensis*) obtained from the 32 sampled horizons.

*Uvigerina peregrina*, which occurs continuously in the section, is a shallow infaunal benthic species whose *δ*^18^Ο and *δ*^13^C values mirror the physical conditions and compositions of ambient bottom waters at the time of calcification. Time-series analysis of *δ*^18^Ο values measured in *Uvigerina peregrina* constitutes a powerful practical tool for global–scale correlation (Elderfield et al., 2012). *Globigerinoides ruber*, which also occurs continuously in the section, is a symbiont bearing planktonic species that preferably calcifies in the oligotrophic upper-most part of the water column (Schiebel and Hemleben, 2017). *Globoconella inflata* occurs in 29 out of the 32 collected samples and is an algal symbiont-barren planktonic species that calcifies mainly at intermediate water depths of the photic zone in stratified water masses or in nutrient-rich surface waters (Schiebel and Hemleben, 2017). In the photosymbiotic species *G. ruber*, there is a positive correlation between shell size and chlorophyll-a content, and the symbiotic abundance of single *G. ruber* is scaled to the growth of the foraminifer (Ezard et al., 2015). This affects the *δ*^13^C composition of the shell through the lifetime of the foraminifera. To limit such an effect, all isotopic measurements on foraminifera were performed on shells handpicked from the 250–355 µm size fraction. About 10 to 15, 20 to 25 and 15 to 20 shells of *U. peregrina*, *G. ruber* and *G. inflata*, respectively, were analyzed per sample to supply sufficient calcium carbonate and to limit the effect of individual variation on the isotopic values (dataset available in Agiadi et al. (2024b). The foraminifera shells analyzed were free of carbonate infilling or visible dissolution features. The samples were ultrasonically cleaned in distilled water for 30 min, and weighed prior to isotopic analyses.

Otoliths are aragonitic, incremental hard parts of the inner ear of teleost fishes, which are metabolically inert (Campana, 1999). Their *δ*^18^Ο values depend on temperature and the *δ*^18^Ο values of ambient water, and they are not affected by fish somatic or otolith growth (Kalish, 1991a). In a paleoceanographic setting, it has also been shown that the *δ*^18^O values of otoliths of epipelagic fish are correlated with those of planktonic foraminifera, whereas the *δ*^18^O values of benthic fish otoliths are correlated with those of benthic foraminifera from the same sediment samples (Agiadi et al., 2024a). The *δ*^13^C of otoliths reflects the *δ*^13^C of the fish diet and that of the dissolved inorganic carbon, and it is therefore a proxy of the fish metabolic rate (Chung et al., 2019a, b; Kalish, 1991a; Martino et al., 2020; Smoliński et al., 2021; Solomon et al., 2006; Trueman et al., 2016, 2023) that has already been successfully applied in fossil otoliths as well (Agiadi et al., 2024a; Wurster and Patterson, 2003).

The fish species *Hygophum benoiti* and *Ceratoscopelus maderensis* were selected because they are among the most common myctophids still living today in the Mediterranean (Mytilineou et al., 2005; Olivar et al., 2012), with very broad geographic distributions (Eduardo et al., 2021; Olivar et al., 2017; Zelck and Klein, 1995), and their otoliths are the most abundant in the Pleistocene fossil record (Agiadi et al., 2010, 2011, 2018, 2023; Girone et al., 2006). The otoliths were sonicated in methanol (MeOH) for about 10 s to remove adhering clay particles, rinsed at least five times in ultraclean water and dried under a clean-air flow. In general, otoliths suffer little from diagenetic alteration, mostly maintaining their microstructure, chemistry and mineralogy, and remain aragonitic even after millions of years (Dufour et al., 2000; Prasanna et al., 2021). Nevertheless, we selected otoliths preserving well all of their morphological characteristics without secondary coloration and/or signs of bioerosion or encrustation that may point toward extensive diagenetic alteration (Agiadi et al., 2022). In addition, randomly selected specimens were observed under a stereoscope to confirm their good preservation state (Agiadi et al., 2022; Antonarakou et al., 2019; Dufour et al., 2000). For the otoliths, a single specimen per sample was crushed, homogenized and analyzed.

Both foraminiferal and otolith *δ*^18^Ο and *δ*^13^C measurements were performed using an Elemental Multiprep system in line with a dual Inlet IsoPrime^TM^ Isotope Ratio Mass Spectrometer (IRMS) at the Université Claude Bernard Lyon 1, France. The principle of the fully automated device is to react the calcium carbonates of foraminifera or otoliths with anhydrous phosphoric acid at 90 °C to generate CO_2_, which is then analyzed with IRMS. Isotopic compositions were quoted in the standard *δ* notation in per mil (‰) relative to V-PDB for both ^18^O/^16^O and ^13^C/^12^C. All sample measurements were adjusted to the international reference NBS18 (*δ*^18^O_V-PDB_ = −23.2‰; *δ* ^13^C _V-PDB_ = −5.01‰) and an in-house Carrara Marble standard (*δ* ^18^O_V-PDB_ = −1.84‰; *δ* ^13^C _V-PDB_ = +2.03‰) calibrated against international references NBS18 and NBS19. With the Carrara marble, the repeatability of the *δ* ^18^O and *δ* ^13^C measurements was 0.16‰ and 0.08‰ (2σ, n = 32), respectively. With NBS18, the repeatability of the *δ* ^18^O and *δ* ^13^C measurements was 0.24‰ and 0.16‰ (2σ, n = 16), respectively. Replicate analyses of 13 foraminifera and 20 otolith samples generated replicate data within the repeatability of the method for both *δ*^18^O and *δ* ^13^C.

### Biomarker analysis

The biomarker analyses followed standard grinding, extraction, purification, and measurements detailed elsewhere (Besiou et al., 2024). Briefly, between 30 and 60 g of all 32 collected samples were dried and powdered. Lipids were extracted using a Soxhlet apparatus with a mixture of dichloromethane (DCM) and MeOH (7.5:1; *v*:*v*) and pre-extracted cellulose filters. All extracts were evaporated to near dryness under N_2_ flow with a TurboVap LV, obtaining the total lipid extracts (TLE). Subsequently, elemental sulfur was removed using 10% HCl-activated and neutralized, and solvent-cleaned Cu shreds. The TLE vials containing the activated Cu and magnetic rods were stirred overnight on a rotary plate. The samples were then filtered over a Na_2_SO_4_ column, dried, and desulfurized again until no reaction with Cu could be observed. A fraction of the TLE was separated into fractions by solid-phase SiO_2_ gel column chromatography based on mixtures of solvents of increasing polarity. The apolar fraction was eluted using a mixture of *n*-hexane and DCM (9:1, *v*:*v*), the ketone fraction using DCM, and the polar fraction using a mixture of DCM/MeOH (1:1, *v*:*v*). The ketone fraction was analyzed using Gas Chromatography Mass Spectrometry (GC-MS), but the samples presented only seldom traces of alkenones, which could not be used for complementary SST reconstruction as initially intended. The polar fraction containing glycerol dialkyl glycerol tetraethers (GDGTs) was dried under N_2_, dissolved in a 1 ml mixture of *n*-hexane (*n*-hexane)/isopropanol (IPA) (99:1, *v*:*v*), dispersed for ∼30 s using an ultrasonic bath, then filtered over a 0.45 mm polytetrafluoroethylene (PTFE) filter using a 1 ml syringe, and subsequently concentrated. The filtered polar fraction was measured using a Shimadzu High-performance liquid chromatography (HPLC), with two Ultra high-performance liquid chromatography (UHPLC) silica columns (2.1 × 150 mm, 1.7 μm) in series, connected to a 2.1 × 5 mm pre-column coupled to an ABSciex 3200 Qtrap chemical ionization mass spectrometer (HPLC/APCIeMS) at Senckenberg Biodiversity and Climate Research Centre, Frankfurt (SBiK-F). The column temperature was 30 °C and the flow rate was 0.2 ml × min^-1^. The GDGTs were eluted isocratically using 18% B for 25 min, then a linear gradient to 35% B in 25 min, followed by a ramp to 100% B in 30 min, where B in *n*-hexane/isopropanol (9:1; *v*:*v*) and A is *n-*hexane. For each sample a 5 µl injection volume was used and GDGT detection was achieved through single ion monitoring for selected scanned masses: 1018, 1020, 1022, 1032, 1034, 1036, 1046, 1048, 1050, 1292, 1296, 1298, 1300, and 1302. The quantification of GDGTs was performed using the Analyst software and the peaks were integrated manually, more than twice for each sample to control for human-induced analytical bias. Because of the absence and/or traces of alkenones we were unable to estimate SST using the U^K^’_37_ paleothermometer. However, we found easily identifiable amounts of isoprenoidal (iso)GDGTs produced mainly by Archaea (Schouten et al., 2013) in open marine settings.

### Sea surface temperature estimation

We applied TEX_86_ paleothermometry (Kim et al., 2010; Schouten et al., 2002) to estimate the sea surface temperature (SST), as follows (Schouten et al., 2002) (Eq. 3):

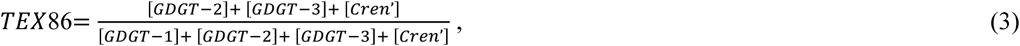

where *GDGT-1*, *GDGT-2, GDGT-3* and *Cren’* are isoprenoid GDGTs. TEX_86_ values were converted into SSTs using the calibration and recommendation of Kim et al. (2010) to apply TEX_86_ ^H^ > 15 °C for regions outside the polar and subpolar latitudes as follows (Eqs. 4 and 5):

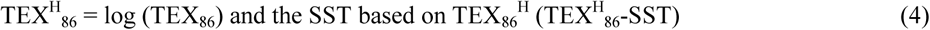

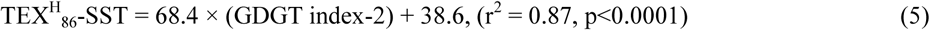

where (Eq. 6):

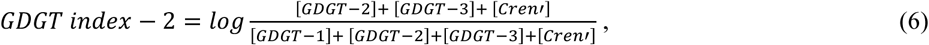

SST calculations based on TEX_86_^H^ have a calibration residual standard (1σ) error of ± 2.5 °C (based on 255 core-top sediments using Kim et al. (2010) calibration equation) (full measurements available as the dataset Agiadi et al. (2024b). The temperature error between sample replicates was ± 0.3 °C. Thus, the propagated error in the temperature estimates is ± 2.52°C.

### Sea surface salinity estimates

Foraminiferal *δ*¹⁸O reflect both the temperature and the ambient oxygen isotopic composition of seawater (*δ*¹⁸O_sw_) in which the shells precipitated. This is true for both planktonic and benthic taxa, but in the case of benthics the *δ*¹⁸O values give insight about the deep water temperatures and sea-level/ice volume changes (Elderfield et al., 2012; Hoogakker et al., 2024). *δ*¹⁸O_sw_, in turn, is influenced by global ice volume and ocean salinity. When the temperature (T) component is accounted for, foraminifera calcite *δ*¹⁸O can be used to estimate past salinity changes, as *δ*¹⁸O_sw_ covaries linearly with sea surface salinity (SSS). Both increase with evaporation and decrease with the addition of low-*δ*¹⁸O freshwater (Legrande and Schmidt, 2006).

Building on this concept and in line with earlier studies (Kontakiotis et al., 2022; Vasiliev et al., 2019), we employed a multi-proxy geochemical approach to estimate regional past SSS variability. To isolate *δ*^18^O_SW_ from the measured foraminiferal *δ*^18^O values, we used *δ*^18^O measurements from *Globigerinoides ruber* shells (*δ*^18^O_rub_), the most surface-dwelling species. The temperature component was removed using the *Orbulina universa* low-light paleotemperature equation (Eq. 7) of Bemis et al. (1998), which is widely regarded as the most accurate for estimating paleo-SST in symbiotic planktonic foraminifera from subtropical settings in deep time (Williams et al., 2005):

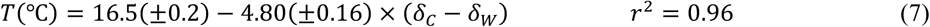

Removing the temperature component from the *δ*^18^O values using the paleotemperature equation enables the estimation of salinity. To account for the effect of continental ice volume, we utilized the Pleistocene eustatic sea-level curve of Rohling et al. (2014). Specifically, we linearly interpolated the sea-level curve to the studied time period and generated a record of global *δ*^18^O_SW_ changes associated with the melting of continental ice sheets during the Early and Middle Pleistocene. This was achieved by converting the sea-level data into mean ocean *δ*^18^O changes and applying a 0.011‰ increase per meter of sea-level drop (Fairbanks and Matthews, 1978; Lea et al., 2002; Schrag et al., 1996). Furthermore, the *δ*^18^O values of *G. ruber* were corrected for species-specific disequilibria from ambient seawater equilibrium (“vital effects”) by subtracting 0.48‰ (Peeters et al., 2002; Pracht et al., 2019). We then subtracted this component from the *δ*^18^O_SW_ profile to derive the regional ice volume-free residual (*δ*^18^O_IVF-SW_). The calculated *δ*^18^O_IVF-SW_ values reflect changes in regional hydrography and are therefore considered to approximate local variations in sea surface salinity (SSS). To convert the *δ*^18^O_rub_ signal into absolute SSS values, we applied the modern *δ*^18^O_SW_–salinity relationship for the Mediterranean Sea of Pierre (1999) that refers to the surface waters with salinities greater than 36.2 (Eq. 8):

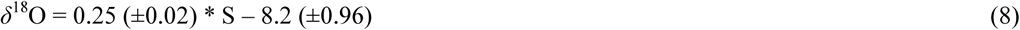

The uncertainty in salinity estimates arises from the propagation of errors associated with temperature estimates, *δ*^18^O_SW_ reconstructions, and the *δ*^18^O_SW_–salinity calibration equation. As demonstrated above, the error in the temperature estimates is 2.5 °C. Based on the *δ*^18^O analytical error and the error associated with the paleotemperature equation, the propagation error for *δ*^18^O is 0.63‰. Using the *δ*^18^O_SW_ – salinity relationship of Pierre (1999), the average propagated uncertainty in salinity reconstruction is calculated to be ∼5.4. In the accompanying graphs, the propagated salinity error was calculated for each sample individually, using the sample-specific *δ*^18^O_SW_ residual.

### Foraminiferal biomasses

Following the method introduced by Schiebel and Movellan (2012) and Movellan et al. (2012), the sediment samples were dry-sieved into seven size categories: < 125 μm, 125–150 μm, 150–200 μm, 200–250 μm, 250–315 μm, 315–400 μm, 400–500 μm, and > 500 μm. The < 125-μm residuals were excluded from the analyses following Milker et al. (2017) and van der Zwaan et al. (1990). We counted the number of planktonic and benthic foraminifera in each size fraction. We only counted individuals whose shell or valve was at least half-preserved. We estimated the planktonic foraminiferal biomass (*pfb*) based on the empirical relationship between test size and protein mass (equivalent of biomass) developed by Schiebel and Movellan (2012) for 26 planktonic foraminiferal species (Eq. 9):

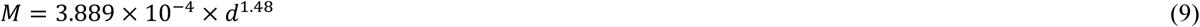

where *M* is the mass of the protein content (μg) and *d* is the minimum diameter of the indivitual (μm). The benthic foraminiferal biomass (*bfb*) was estimated using the relationship of Movellan et al. (2012) for *Ammonia tepida* (Eq. 10), which is currently the only available equation for estimating benthic foraminiferal biomass:

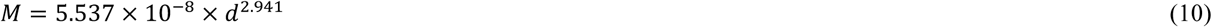

The total foraminifera biomasses (*fb*), *pfb*, and *bfb* (total and for each size class) were then calculated as the *M* times the number of tests counted in the sample or fraction divided by the dry weight of the sediment sample in g.

### Ostracod and sponge spicule size abundances and distributions

Ostracod and sponge spicule abundances and size distributions were obtained in order to detect and evaluate changes in secondary production, which would affect overall ecosystem productivity and therefore nutrient availability for mesopelagic fishes, throughout the study time interval. Ostracod and sponge spicules were counted at each sediment fraction and reported here as the number of valves or spicule fragments, respectively, per gram of sediment. Ostracods are very sensitive to changes in the oxygen content of seawater (Huang et al., 2019), and Pleistocene paleoclimatic changes severely affected the ostracod assemblage composition and abundance in many areas worldwide (Alvarez Zarikian, 2016; Alvarez Zarikian et al., 2022; Cronin et al., 2014; DeNinno et al., 2015; Huang et al., 2019), including the Mediterranean (Rossi et al., 2018).

Sponges are very attuned to water depth, pH, temperature, and oxygenation, making their assemblages and abundance very good indicators of paleoenvironmental conditions (Łukowiak, 2020). Spicule abundance in the sediment is related to the abundance of sponge individuals (Łukowiak, 2020), reflecting biogenic silica accumulation, and has been used as a proxy of sea bottom conditions (Alvarez et al., 2017).

### Statistical analyses

To evaluate the effect that paleoenvironmental changes had on mesopelagic fishes, we first needed to establish when these changes took place, in the study area. We used the Sequential T-test Analysis of Regime Shifts (STARS) algorithm (Rodionov, 2004) to detect rapid, statistically significant shifts in each paleoenvironmental parameter (SSS, SST, productivity) separately. This algorithm uses the t-test to determine whether sequential records in a time series represent statistically significant departures from the mean value observed during the preceding period. The algorithm outputs a value of the regime shift index (RSI), which represents the sum of the normalized anomalies indicating the shift magnitude. The STARS algorithm is able of statistically detect significant shifts even at the edges of a time series, which is why it has been the preferred statistical method in many previous studies (e.g., (Agiadi et al., 2024a; Harning et al., 2021; Lu et al., 2025; Yuan et al., 2020). Regime shifts are rapid reorganizations of the ecosystem that are reflected in the analysed parameter. Therefore, the cause may be different in each case depending on the analysed parameter. To detect regime shifts, we first checked the time series of all the variables measured for normality using the Shapiro-Wilk test. For the variables whose values did not exhibit a normal distribution, we applied the L-method, which is based on the Mann-Whitney U test (Lanzante, 1996). For the variables whose values were normally distributed, we used the STARS algorithm, setting Huber’s weight parameter to 1 and the window size arbitrarily to 10. For both the L- and the Rodionov’s methods, the *p*-value was set at 0.05.

Correlation testing was performed to detect covariance between temperature or productivity, the isotopic signals in the fossil skeletal material of the different organisms and their biomasses or abundances. Specifically, we tested correlation between: a) SST and all the *δ*^18^Ο values; b) TOC (as a proxy of productivity and preservation of organic matter) and all the *δ*^13^C values; c) SST and foraminifera biomasses (*fb*, *pfb*, *bfb*); d) TOC and the *δ*^13^C of foraminifera, against the foraminiferal biomasses; e) SST and the ostracod weighted average body size; and f) TOC and the foraminiferal *δ*^13^C, against the ostracod body size. In all cases, we used the Spearman rank correlation coefficient (95% confidence level).

All analyses were performed in R (version 4.3.3) (R Development Core Team, 2024) with the package *rshift* (Room et al., 2020).

### Lifetime-average depth of mesopelagic fish

Mesopelagic fishes live in the water column, migrating between the mesopelagic and the euphotic zones during the DVM. The precise time spent in each zone depends on the species, the season and the ontogenetic stage. Here, we estimate the average water depth the fish lived in throughout its lifetime, which we term lifetime-average depth, for each species and at each time interval (height along the studied section). Because otoliths grow throughout the lifetime of the fish, and because they biomineralize day and night, their δ¹⁸O values are actually lifetime-averages (Campana, 1999). Consequently, the depths obtained from the model are also fish lifetime-average depths.

The obtained foraminiferal *δ*^18^O data representing the thermohaline gradient was used to estimate the position of the fishes in the water column. An exponential model was fitted using a log-linear regression on the *δ*^18^O values of foraminifera living at different water depths to infer lifetime-average depths in the water column where the two mesopelagic fish species, *Ceratoscopelus maderensis* and *Hygophum benoiti*, were living (fish lifetime-average depths) during each individual climatic cycle. The benthic foraminifera *Uvigerina peregrina* grew at the sea bottom, and therefore its living depth used in our model was the paleodepth estimated in each sample based on the *%P/(B+P)* ratio. We assigned a living depth distribution of 100–300 m to *Globoconella inflata*, because this deep-dwelling planktonic foraminifera is consistently most abundant at these cool and nutrient-rich subsurface water depths (Rohling et al., 2004; Schiebel and Hemleben, 2017). We finally assigned a depth distribution of 10–50 m to *Globigerinoides ruber*, since this surface-dwelling photosymbiotic species is most abundant at warmer and oligotrophic surface mixed-layer depths (Rohling et al., 2004; Schiebel and Hemleben, 2017). Since direct measurements of the exact depth (*y*) where planktonic foraminifera were actually living could not be obtained, we simulated the values within the ecological depth ranges of each species. For each species, depths were drawn from a uniform distribution over the respective depth ranges, considering equal probability across the range.

We fitted an exponential model using a log-linear regression where the realizations of our response variable *y*_i_ represent the observed depths for each foraminiferal species (ecological depth ranges). The response variable was log-transformed during modeling to establish an exponential relationship with *δ*^18^O (*x*) (Eq. 11):

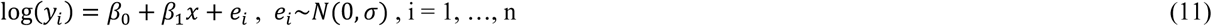

where we assumed that log(*y*_i_) follows a normal distribution and that the linear predictor is made of an intercept *β*_0_ and the relationship with *δ*^18^O. Separate models were fitted for glacials (MIS 22, 20 and 18) and interglacials (MIS 23, 21 and 19) to account for potential differences in the *δ*^18^O-depth distribution relationships between climate cycles.

Using the model at each individual MIS, we thus estimated the fish lifetime-average depths based on the otolith *δ*^18^O values. This model approach assumes that the thermal gradient remains the same within each MIS and that the water column is always well mixed. Although this is a simplified model of thermohaline conditions, the outputs offer a valuable perspective on the evolution of the lifetime-average depth and DVM patterns of mesopelagic fishes. The code for our model is available in (Fuster-Alonso et al., 2024).

## Results

### Paleodepths

The *%P/(P+B)* ratio ranges between 45% and 76%, and the 32 collected samples consequently deposited at the outer shelf (5 samples) and upper bathyal (27 samples) paleodepths, with estimates ranging between ∼176 m and ∼528 m. Although absolute paleodepth values based on *%P(P+B)* need to be taken cautiously (Van Hinsbergen et al., 2007; van der Zwaan et al., 1990), three main intervals with lower approximate values around ∼200 m are recorded in the lower part of the section at heights between 40 and 60 cm, in the middle part of the section between 170 and 180 cm, and in the uppermost part of the studied deposits above 270 cm (Fig. 1). Overall, the highest upper bathyal paleodepths are reached at three main intervals: below the height of 40 cm, between 70 and 160 cm, and between 190 and 270 cm.

### Oxygen and carbon isotopic compositions

The *δ*^18^O values of bulk sediments (*δ*^18^O_bulk_) vary between −2.82 (LR30) and −0.70 ‰ (LR16) (full measurements available as the dataset Agiadi et al. (2024b). They are relatively higher overall up to height 160 cm, and lower thereafter. The *δ*^13^C_bulk_ values range from −2.68 (LR25) to −0.96 ‰ (LR12). Lower values appear in general until LR16 and higher thereafter.

The *δ*^18^O_ruber_ values range from −0.62 (LR08) to 1.22 ‰ (LR01), *δ*^18^O_inflata_ from 1.23 (LR17) to 2.19 ‰ (LR01), and *δ*^18^O_peregrina_ from 1.51 (LR22) to 2.56 ‰ (LR30). The three species of foraminifera exhibit *δ*^18^O values that reflect their depth-related ecology, with highest values in the benthic species *Uvigerina peregrina*, and lowest *δ*^18^O values in the surface mixed-layer species *Globigerinoides ruber* (Fig. 2). *Globoconella inflata* exhibits intermediate *δ*^18^O values that are closer to those of *U. peregrina* (0.5 ‰ average gradient) than to those of *G. ruber* (1.7 ‰ average gradient), then confirming that *G. inflata* mineralized at cooler/deeper water depths in the water column than *G. ruber*. *Globigerinoides ruber* and *U. peregrina* exhibit an average gradient of 1.9 ‰ between surface and bottom waters. *Uvigerina peregrina*, *G. inflata* and *G. ruber* show similar overall variation trends throughout the section, but with discrepancies notably recorded between *G. ruber* and the two other species.

**Figure 2.**
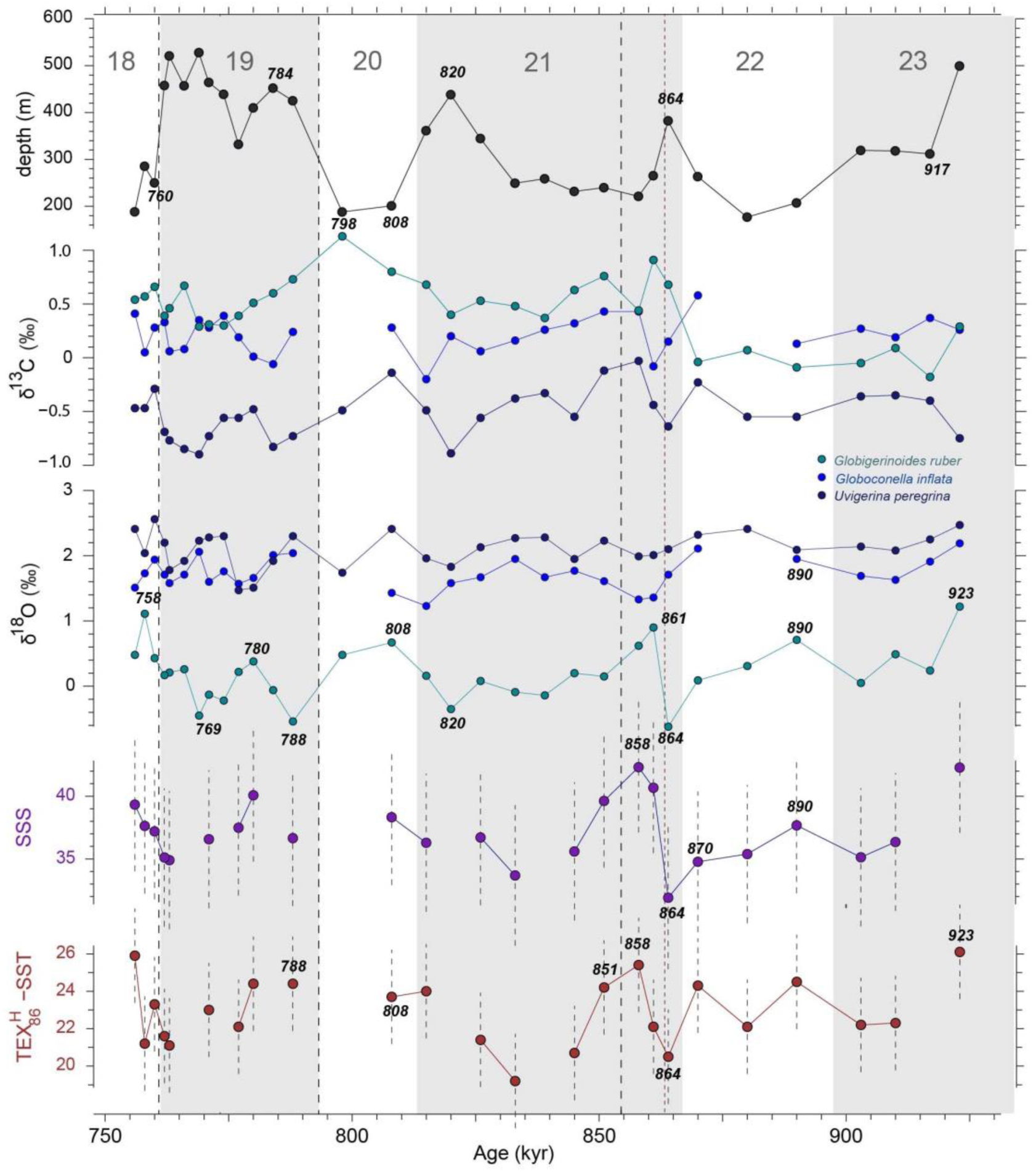
Paleoclimatic and paleoceanographic conditions at Lardos. Paleodepth estimates based on the *%P/(B+P)* ratio. *δ*^18^O and *δ*^13^C values of the surface-dwelling planktonic foraminifera *Globigerinoides ruber*, the deeper-dwelling planktonic foraminifera *Globoconella inflata*, and the benthic foraminifera *Uvigerina peregrina*. TEX_86_-derived sea surface temperature (SST) and salinity (SSS) data, where bars are used to visualize the ±2.5 °C error. The three red wide-dashed vertical lines indicate the three regime shifts identified using the STARS algorithm (total organic carbon (TOC), *δ*^18^Ο_peregrina_ and *δ*^18^Ο_ruber_ at 760 ka B.P.; *δ*^13^C_peregrina_ and the ostracod weighted average body size at 788 ka B.P.; and *δ*^13^C_ruber_ at 851 ka B.P.), whereas the narrow-dashed line shows the regime shift identified using the L-method (ostracod weighted average body size at 864 ka B.P.). Grey-shaded areas represent interglacials, whereas white areas are glacials; the number of each MIS is indicated on top. Italicized numbers show the ages corresponding to each data point.

For the fishes, *δ*^18^O_benoiti_ values range from 2.56 (LR28) to 7.35 ‰ (LR02), and *δ*^18^O_mader_ ranges from −3.3 (LR16) to 3.50 ‰ (LR10). There is no significant correlation between otolith weight (see full dataset in (Agiadi et al., 2024b)) and *δ*^18^O. *δ*^13^C_ruber_ values range from −0.18 (LR02) to 1.13 ‰ (LR19), with lower values until LR07.

*δ*^13^C_inflata_ values range from −0.2 (LR17) to 0.58 ‰ (LR07), and *δ*^13^C_peregrina_ ranges from −0.89 (LR16) to −0.3 ‰ (LR10). Ostracod *δ*^13^C_baird_ values, only measured in the upper part of the section, ranges from −1.18 (LR28) to −0.61 ‰ (LR25). For the fishes, *δ*^13^C_benoiti_ ranges from −5.25 (LR32) to 0.99 ‰ (LR09), with generally more negative values in the upper part of the section, and *δ*^13^C_mader_ range from −6.70 (LR07) to 2.92 ‰ (LR16). The distributions of the *δ*^13^C_mader_ and *δ*^13^C_benoiti_ values are shown in Fig. S1. There is a significant correlation between otolith weight and otolith *δ*^13^C for both *Hygophum benoiti* (*rho* = 0.45, *p* = 0.04) and *Ceratoscopelus maderensis* (*rho* = 0.80, *p* = 0.0003). The different foraminifera and fish species show distinctive patterns of *δ*^13^C and *δ*^18^O (Fig. S2).

### Revised age model

Our results reveal that the lowermost part of the Lardos section recovers the Early/Middle Pleistocene boundary (774 ka B.P.) and the interval between MIS 23 (*partim*) and MIS 18 (Fig. 1). By digging a pit at the base of the section originally studied by Titschack et al. (2013), we have extended the record covered by the Lardos section down to MIS 23. At heights 30 to 40 cm (samples LR04–05), the *δ*^18^O values of all three species of foraminifera increase and the paleodepth shows a sharp decrease. Such changes are proposed to correlate with those occurring in the global (Lisiecki and Raymo, 2005a) and Mediterranean (Konijnendijk et al., 2015) benthic *δ*^18^O stacks, and with that occurring in the planktonic foraminifera Mediterranean *δ*^18^O Medstack (Wang et al., 2010) between MIS 23 and MIS 22 (Fig. 1). In this re-examination of the age model of the Lardos section, we assign the MIS 22/21 termination (866 ka B.P.; Lisiecki and Raymo, 2005) to the height of 65 cm, in accordance with Titschack et al. (2013). Such a level, characterized by a change in the slope in all foraminiferal *δ*^18^O records and a sharp increase in paleodepth, is proposed in agreement with those occurring in open-ocean reference records (Fig. 1). We also delineate the MIS 21/20 (814 ka B.P.) and MIS 20/19 (790 ka B.P.) terminations to the heights 165 (LR17–18) and 185 cm (LR19–20), respectively. Within the sedimentary interval that we allocate to MIS 20, and that shows shallower paleodepths, both *U. peregrina* and *G. ruber* exhibit increasing *δ*^18^O values similar to those of open-ocean reference records (Fig. 1). Finally, we assign the MIS 19/18 termination (761 ka B.P.) to the height of 275 cm (LR29–30), in accordance with the previously published chronostratigraphy (Titschack et al., 2013). Such a level shows increasing *δ*^18^O values in all foraminiferal taxa, similar to the open-ocean records, and a sharp decrease in paleodepth.

Based on these tie-points, we estimate that deposition in the studied hemipelagic sediments of the Lardos section occurred with an average sedimentation rate of ∼1.8 cm/kyr, a value that was linearly extrapolated from the MIS 23 and MIS 18 intervals. Sedimentation rates reached lower average values of 1.0 and 1.1 cm/kyr in glacial intervals (MIS 22 and MIS 20, respectively), and higher average values of 1.8 and 2.8 cm/kyr in interglacial intervals (MIS 21 and MIS 19, respectively). The greater average value obtained from MIS 19 partly results from the ∼25 cm-thick debris flow horizon cropping out at 2.3 m height. According to this refined age model, the samples analyzed in this study were deposited within the MIS 23–18 interval between ∼923 and ∼756 ka B.P. (Fig. 1).

### Organic content in the sediments

The BIT index is lower in the older part of the section (i.e., lower input of terrestrial organic matter), with a minimum of 0.45 at 870 ka B.P. (Fig. 3). It then increases (toward more terrestrial organic matter input) to reach a maximum of 0.80 at 815–808 ka and remains high in the younger part of the section. TOC ranges from 0.152 to 0.266% and TN from 0.027 to 0.042%, making the C/N ratio range from 5.14 to 7.27 (Fig. 3). From 923 ka B.P., both TOC and TN increase to reach 0.266% and 0.042%, respectively, at 864 ka B.P.. In the interval 861–820 ka B.P., they show relatively high values, which then decrease to a minimum at 798 ka B.P. of 0.180 and 0.032%, respectively. Intermediate values appear from 788 to 762 ka, and then low values for both TOC and TN until 756 ka B.P.

**Figure 3.**
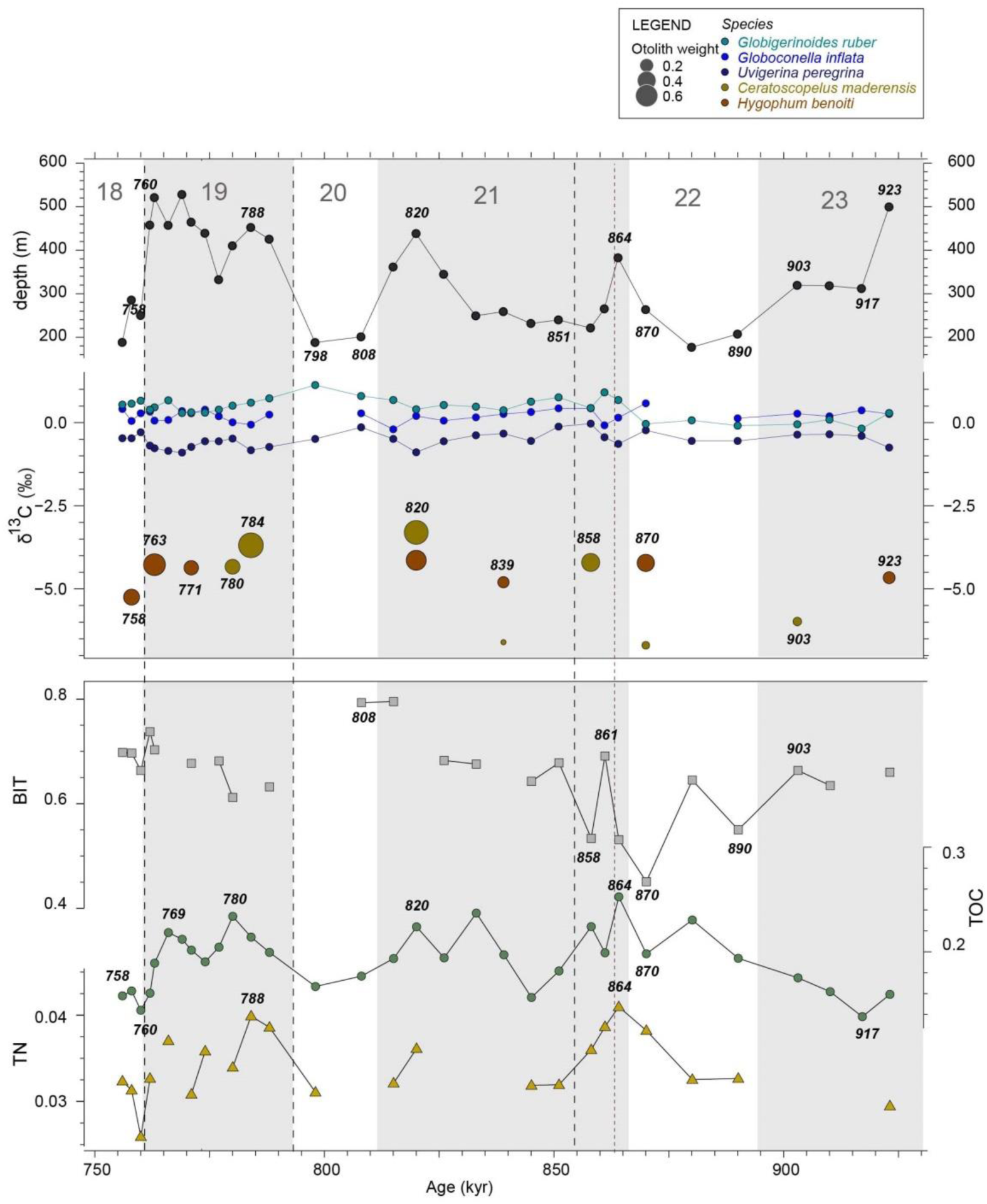
Organic matter and productivity proxy records at Lardos. Paleodepth estimates based on the *%P/(B+P)* ratio. *δ*^13^C values of the surface-dwelling planktonic foraminifera *Globigerinoides ruber*, the deeper-dwelling planktonic foraminfera *Globoconella inflata*, and the benthic foraminifera *Uvigerina peregrina*. The *δ*^13^C of the fish otoliths reflects the *δ*^13^C of the dissolved inorganic carbon in the ambient water (indicated here at different heights along the Lardos section by the foraminifera *δ*^13^C values) and the *δ*^13^C of their diet items. Ultimately, the *δ*^13^C of otoliths is a proxy of the metabolic rate of the fish. The total organic carbon (TOC) and total nitrogen (TN) contents reflect productivity, and the Branched Isoprenoid Tetraether (BIT) index reflects the source of the organic matter in the sediment (which increases with terrestrial influx). Grey-shaded areas represent interglacials, whereas white areas are glacials; the number of each MIS is indicated on top. Italicized numbers show the ages corresponding to each data point.

### Sea surface temperatures and sea surface salinities

SST ranges from 19.2 to 26.1 °C (Fig. 2). From 923 to 864 ka B.P., SST decreases to 20.5 °C. The interval 861–858 ka exhibits increased temperature, but afterwards it is low until 820 ka B.P., when it starts to increase to a maximum of 24.4 °C around 798 ka B.P.. Between 788 and 762 ka B.P., SST shows intermediate, but rather stable values fluctuating around 23°C, and then increases until 756 ka B.P. to 25.9 °C.

SSS ranges from 31.9 to 42.3 (Fig. 2). From 923 to 864 ka B.P., SSS decreases to 31.9, but 861–858 ka B.P. experiences the highest SSS values. Afterwards, SSS decreases again and shows intermediate values after 760 ka B.P.

### Foraminiferal biomass

The foraminiferal biomass is mostly composed of plankton (*pfb*), which is much greater than benthos (*bfb*) at all levels: the minimum is found at 756 ka B.P. (88.3 µg/g for *fb*, 58.8 µg/g for *pfb*, and 29.4 µg/g for *bfb*), whereas the maximum biomass is found at 766 ka B.P. (1763.3 µg/g for *fb*, 1518.2 µg/g for *pfb*, and 245.1 µg/g for *bfb*) (Fig. 4). For both *pfb* and *bfb*, the contributions of the size classes to the total biomass remains proportional (Figs. S3 and S4). From 923 to 880 ka B.P., both *pfb* and *bfb* are low, and then they increase to peak at 861 ka B.P. (867.7 µg/g and 240.4 µg/g, respectively). They remain high until 815 ka B.P., and then decrease until 788 ka B.P. (minima are 303.5 µg/g at 798 ka and 60.7 µg/g at 788 ka, respectively). In general, *bfb* covaries with *pfb*, with two exceptions, at 774 and 769 ka B.P., where *bfb* is particularly low (77.6 and 66.5 µg/g).

**Figure 4.**
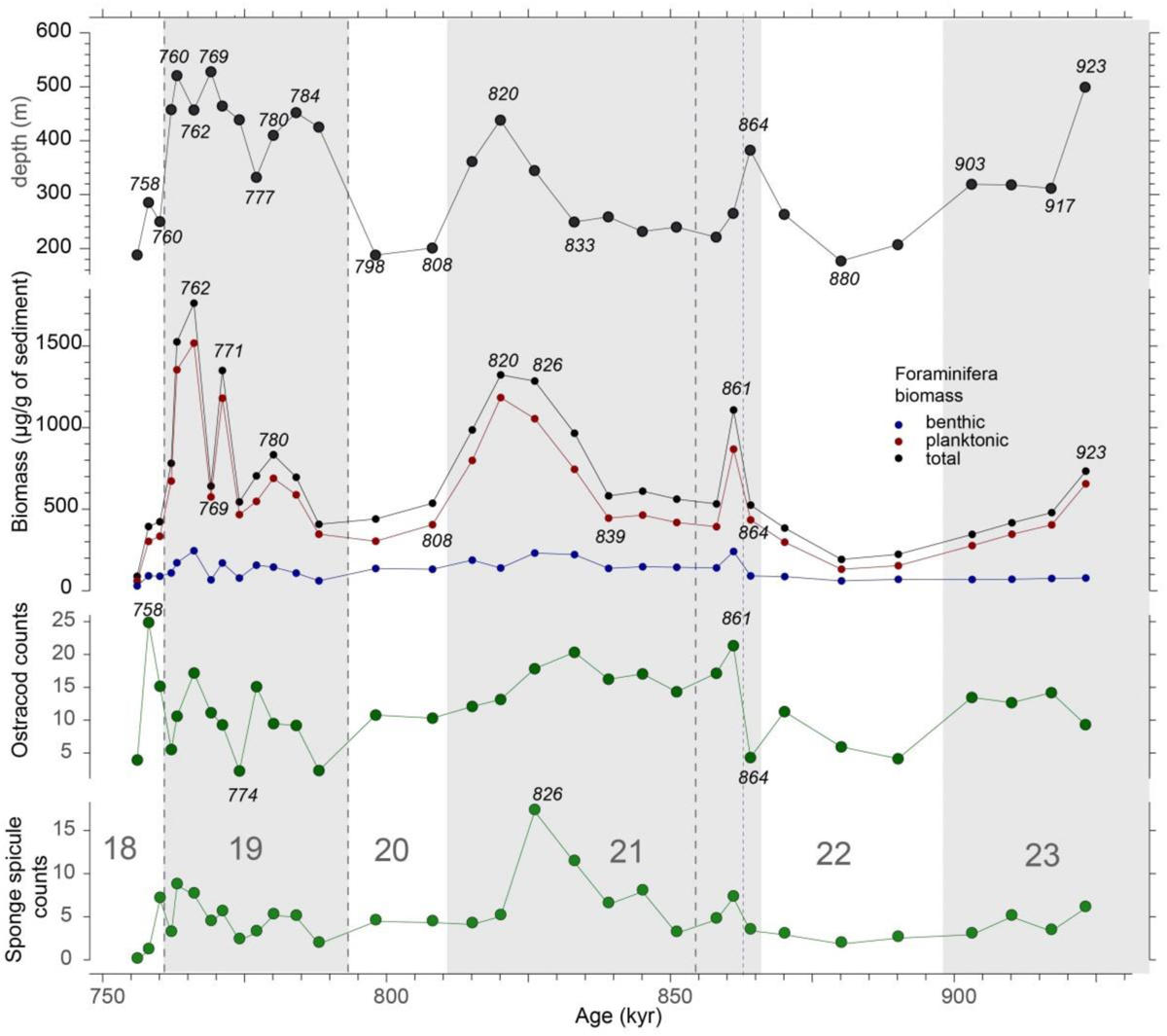
Planktonic, benthic and total foraminiferal biomasses, ostracod and sponge spicule counts at Lardos. Paleodepth estimates based on the *%P/(B+P)* ratio. Grey-shaded areas represent interglacials, whereas white areas are glacials; the number of each MIS is indicated on top. Italicized numbers show the ages corresponding to each data point.

### Ostracod and sponge spicule abundances and size distributions

The total ostracod abundance ranges from 2 valves/g at 774 ka to 25 valves/g at 758 ka B.P. (Fig. 4). From 923 to 903 ka B.P., the medium–high abundance is driven by ostracod valves in the 150–200 μm size-class. The overall number of valves then decreases, and high values are found again from 861 to 820 ka B.P., when their abundance increases across size classes. The abundance after that point fluctuates between low and medium levels, and only shows maximum values at 758 ka B.P. due to highs in the >500, 315–400, and <200 μm size-classes.

The number of sponge spicule fragments per gram of sediment ranges from 2 at 880 ka to 18 at 826 ka B.P. (Fig. 4). The weighted average size of the ostracod valves ranges from 159 μm at 903 ka to 241 μm at 774 ka B.P.. Broadly, similar to ostracod abundance and foraminiferal biomass, the values are generally relatively high around 861–820 ka B.P.. The weighted average size of the spicules ranges from 149 μm at 760 ka to 318 μm at 758 ka B.P.

### Fish lifetime-average depth in the water column

Our model shows a steeper *δ*^18^O_sw_ gradient during interglacials than glacials, predicting greater lifetime-average depth distributions for fish (Fig. 5). Based on the model results, the fish lifetime-average depths range from 474 m at 870 ka to 933 m at 839 ka for *Hygophum benoiti* and from 330 m at 903 ka to 1502 m at 858 ka for *Ceratoscopelus maderensis* (Fig. 6). We could not detect a significant correlation between the modeled fish lifetime-average depth and otolith weight, which suggests that fish lifetime-average depth is not controlled by life history in the same way throughout the studied interval for either species. Additionally, we did not find any significant correlation between the modeled fish lifetime-average depth and SST, total organic carbon (TOC), or *δ*^13^C_bulk_ (full data available as the dataset Agiadi et al. (2024b).

**Figure 5.**
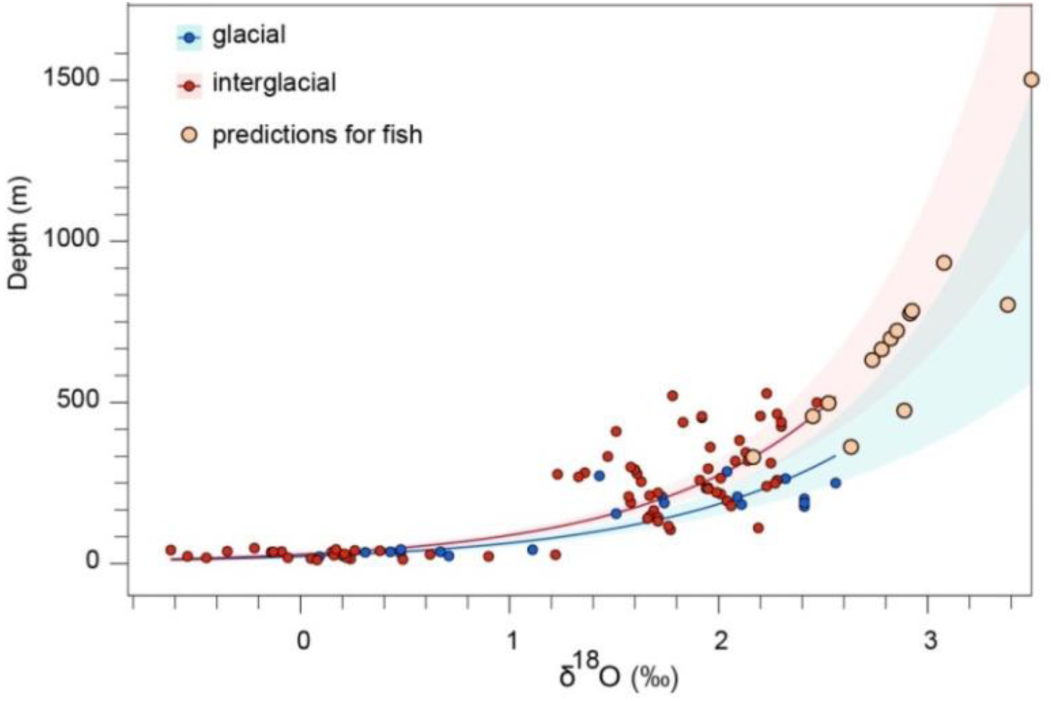
Fitted regression for glacial and interglacial climate cycles with extrapolated fish predictions and 95% confidence intervals.

**Figure 6.**
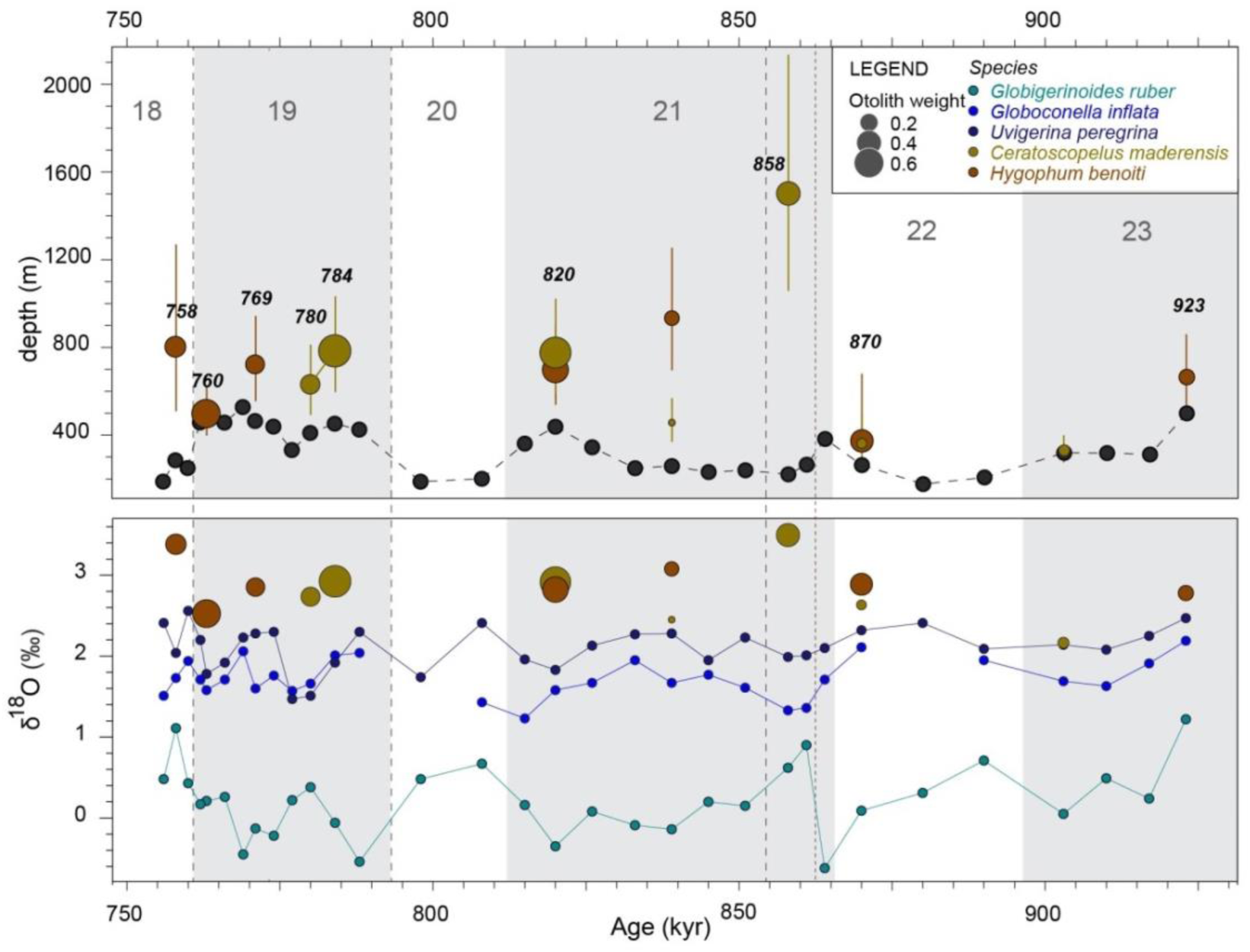
Paleodepth and modeled fish lifetime-average depths where the two mesopelagic fish species lived in the study area, obtained from the *δ*^18^Ο values in their otoliths by modeling the thermohaline gradient using the *δ*^18^Ο values of the different species of foraminifera. The circle diameter scales with otolith weight, reflecting fish size, whereas the vertical lines indicate the fish lifetime-average depth ranges. Grey-shaded areas represent interglacials, whereas white areas are glacials; the number of each MIS is indicated on top. Italicized numbers show the ages corresponding to each data point.

### Regime shifts and covariance

Positive values of the Regime Shift Index (RSI) indicate the presence of regime shifts (95% confidence level) for: a) TOC, *δ*^18^Ο_peregrina_ and *δ*^18^Ο_ruber_ at 760 ka (RSI = 0.940, 1.437 and 0.265, respectively); b) *δ*^13^C_peregrina_ and the ostracod weighted-average body size at 788 ka (RSI = 0.583 and 0.231, respectively); and c) *δ*^13^C_ruber_ at 851 ka (RSI = 0.788). Additionally, a regime shift was found using the Lanzante method for the sponge weighted average size at 864 ka (p = 0.006).

Regarding the foraminiferal biomasses, a medium correlation is detected between: a) SST and benthic foraminifera biomass (*rho* = −0.44; *p* = 0.03); b) *δ*^13^C_ruber_ and benthic foraminifera biomass (*rho* = 0.45; *p* = 0.01); c) *δ*^13^C_inflata_ and total foraminiferal biomass (*rho* = −0.42; *p* = 0.02), planktonic foraminiferal biomass (*rho* = −0.45; *p* = 0.014), and the benthic foraminiferal biomass (*rho* = −0.44; *p* = 0.014); d) *δ*^13^C_peregrina_ and the total foraminiferal biomass (*rho* = −0.47; *p* = 0.007) and the planktonic foraminiferal biomass (*rho* = −0.53; *p* = 0.002); and e) *δ*^13^C_bulk_ and the total foraminifera biomass (*rho* = 0.52; *p* = 0.003), the planktonic foraminiferal biomass (*rho* = 0.52; *p* = 0.003), and the benthic foraminiferal biomass (*rho* = 0.42; *p* = 0.019). Medium correlation is also found between *δ*^13^C_peregrina_ and the ostracod weighted average body size (*rho* = 0.40; *p* = 0.02).

## Discussion

The unique SST reconstruction for the Eastern Mediterranean that is presented here does not march the resolution of the time-equivalent from deep-sea cores in the Western Mediterranean and the Atlantic due to the large differences in sedimentation rates between sites, highlighting the importance of obtaining SST records from deep-sea drilling sites in the Eastern Mediterranean. SST in the study area were consistently warmer than those in the Central and Western Mediterranean, with higher mean SST values found in all climate cycles, between MIS 23 and 18 (Table S1; Fig. 7); this pattern is also observed today (Minnett et al., 2019). The glacial–interglacial SST amplitude is similar (∼5 °C) at both Site LC07 (obtained at water depth of 488 m (Martínez-Dios et al., 2021); Fig. 1) and Lardos, but very different from that recorded at the Atlantic Site U1385 (Fig. 7). The two-fold smaller temperature changes observed in the Mediterranean indicate that the basin responded to the global climate change during the Pleistocene, but the recorded temperatures must have also been influenced by the semi-enclosed configuration of the Mediterranean Sea. SST values at Lardos are also higher than those reconstructed from Site LC07. Part of this difference can be explained by a system similar to the present-day one, where the Eastern Mediterranean is ∼2 °C warmer than the Central Mediterranean (Minnett et al., 2019). Another part of the difference can also be explained by the different proxies used to obtain the SST estimates, i.e. TEX_86_ for Lardos and U*^K^*_37_ for Site LC07. TEX_86_ generally overestimates SSTs by 2–6 °C (Kim et al., 2015), whereas U*^K^*_37_ tends to underestimate SSTs by 2–4 °C (Tierney and Tingley, 2018). Importantly, based on the TOC content of the analyzed samples, we did not capture any of the sapropels (Martínez-Dios et al., 2021) in the study area.

**Figure 7.**
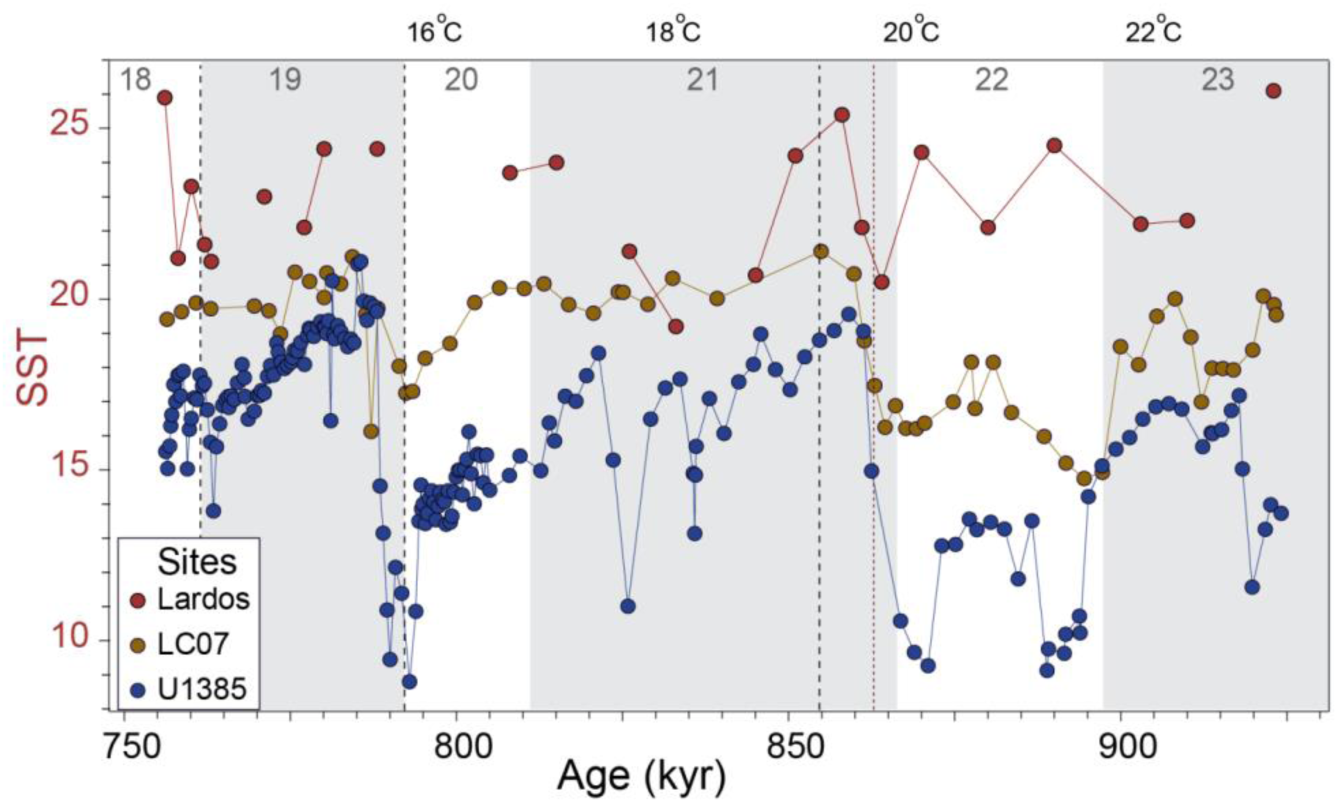
SST time series at: Lardos (this study), Site LC07 (Martínez-Dios et al., 2021) and Site U1385 (Rodrigues et al., 2017). The mean SSTs for all the localities are available as the dataset Agiadi et al. (2024b).

The thermohaline gradient is inferred from *δ*^18^Ο_ruber_ (corresponding to surface waters), *δ*^18^Ο_inflata_ (intermediate waters), and *δ*^18^Ο_peregrina_ (bottom waters) values, considering the paleodepth in the study area (Fig. 2). The gradient between these values is steeper in MIS 23–MIS 22 but attenuates in MIS 21. In the late part of MIS 21, a short warmer period occurs around 820 ka (Fig. 2), but the differences between the foraminifera *δ*^18^Ο values are maintained despite a deeper paleodepth, suggesting that the thermohaline gradient is less steep than before. During MIS 20 and MIS 18, the surface-to-deep sea *δ*^18^Ο gradient is weaker but this can be explained by the shallower sea-levels during glacials (Fig. 2). Our model produces steeper thermohaline gradients during interglacials than during glacials (Fig. 5). This is expected due to the lower thermal conductivity of water masses. As surface water heats up during warm periods, water at depth remains cooler.

Foraminifera are very sensitive to parameters of ambient seawater, such as food supply, light intensity, and sea temperature, which control community composition and pelagic biomass production (e.g., (Gooday and Jorissen, 2012; Schiebel et al., 2018). In the case of the Pleistocene, no consistent responses of foraminiferal biomasses and isotopic compositions have been found on a global scale, and identified productivity impacts have been mostly related to regional changes in nutrient supply through variations in the hydrographic regime (Diester-Haass et al., 2018). With the production of their calcareous shells, both planktonic and benthic foraminifera actively contribute to the biological carbon pump (Grigoratou et al., 2022; Schiebel, 2002). However, quantifying their contribution to carbon export in the geological past has been hampered by the lack of estimates of their biomass. Although foraminifera have long been shown to constitute very powerful tools for paleoceanographic and paleoclimatic reconstructions, their use has been based almost exclusively on the analysis of their assemblage composition, species abundance, and/or elemental and isotopic composition, whereas very little is known about the fluctuations in foraminifera biomass in response to changes in environmental parameters. Planktonic foraminifera biomass is mainly controlled by temperature during the early stage of their development, but food availability becomes more important for mature forms (Grigoratou et al., 2019), whereas benthic foraminiferal biomass is tightly linked to organic matter fluxes and oxygenation (Milker et al., 2019).

Planktonic and benthic foraminiferal biomass changes closely follow the paleodepth changes recorded in the study area (Fig. 4), but we do not anticipate greater accumulation of shells on the seabed due to transport. Since paleodepths exceed the euphotic zone at all times, transport alone would not be expected to increase foraminiferal biomass. Additionally, higher foraminiferal biomass is observed in interglacials than in glacials, and this is confirmed by the ostracod and sponge abundances (Fig. 4). Additionally, planktonic foraminiferal shells in MIS 21 and MIS 19 interglacials are larger overall, with the > 500 μm size fraction increasing in those intervals. Larger shell sizes in planktonic foraminifera are typically associated with temperature and/or primary productivity conditions that are close to the optimum values for these organisms (Schmidt et al., 2004). Therefore, we propose overall higher secondary production during interglacials.

Otolith *δ*^13^C in conjunction with foraminiferal *δ*^13^C are informative of how the metabolic rate of fishes changes through time (Agiadi et al., 2024a). In this case, no important changes in the otolith *δ*^13^C values of the two fish species are observed over time. An exception is the slight drop in *δ*^13^C_benoiti_ in MIS 18, which only occurred from one point. In general, a lower *δ*^13^C_oto_ reflects a higher metabolic rate or a change to higher trophic-level (and therefore lower *δ*^13^C) food items (Chung et al., 2019b). Here, overall, the larger otolith specimens exhibit higher *δ*^13^C values, suggesting that, as the animals grew, they consumed fewer and/or switched to lower-trophic–level items. A possible explanation could be that younger individuals consumed much, primarily detritus, whereas adults became more selective, consuming mesozooplankton.

In this study, we used otolith *δ*^18^Ο as a proxy of mesopelagic fish DVM patterns, since it is a function of seawater temperature and salinity, and it varies according to the lifetime movements of the targeted fish (both vertical along the water column and lateral migrations) (Kalish, 1991b, a). The otolith *δ*^18^Ο values of both mesopelagic fish species are consistently greater than those of foraminifera, which indicates that these fishes experienced much cooler and/or saltier seawater temperatures on average. Both *Ceratoscopelus maderensis* and *Hygophum benoiti* perform DVM today (Backus et al., 1968; Linkowski, 1996). The fact that *δ*^18^Ο_benoiti_ and *δ*^18^Ο_mader_ fluctuate between much more positive and almost the same values of *δ*^18^Ο_peregrina_ suggests that they spent only part of their day in the study location, migrating from greater depths in the water column. Furthermore, with the present data, the *δ*^18^Ο_benoiti_ and *δ*^18^Ο_mader_ variability cannot be related to the *δ*^18^Ο_foram_ variability, nor the SST changes. Particularly, for *Ceratoscopelus maderensis*, this variability shows high amplitude, suggesting that this species changed its DVM pattern and/or its lifetime-average depth within the studied time interval. Today, the common, extant species *Ceratoscopelus maderensis* shows different depth distribution ranges in the Atlantic Ocean (Badcock and Merrett, 1976; Killen et al., 2010; Kinzer and Schulz, 1985) than in the Mediterranean Sea, and even within the latter (up to 1480 m in the Levantine Sea (Goren and Galil, 2015), but up to 2500 m in the Ionian Sea (Olivar et al., 2012). Our *δ*^18^Ο–depth model estimates of the lifetime-average depth of the two mesopelagic fishes show high variability.

In MIS 23–22, *Ceratoscopelus maderensis* (indicated by small otolith specimens corresponding to young individuals) are distributed on average at lifetime-average depths of 330 m (Fig. 6), which is at the very shallow end of the present-day distribution of this species, suggesting that the individuals (at least the young ones), as observed today in the Mediterranean (Olivar et al., 2012), were performing DVM and/or occupying shallower depths in the water column (Fig. 8). The modeled lifetime-average depths of *Hygophum benoiti* are greater in MIS 23 than modern ones, but the same in MIS 22. However, we cannot interpret these results as absence of DVM in MIS 23, because we have only examined a few specimens per species as replicates. A greater number of otolith specimens should be analyzed per sampling unit to reconstruct the frequency distribution of the lifetime-average depths of each species to be able to confidently exclude DVM. Large adults of both species show shallower lifetime-average depths (776 and 698 m, respectively) in the later part of MIS 21, and the same pattern is observed in MIS 19.

**Figure 8.**
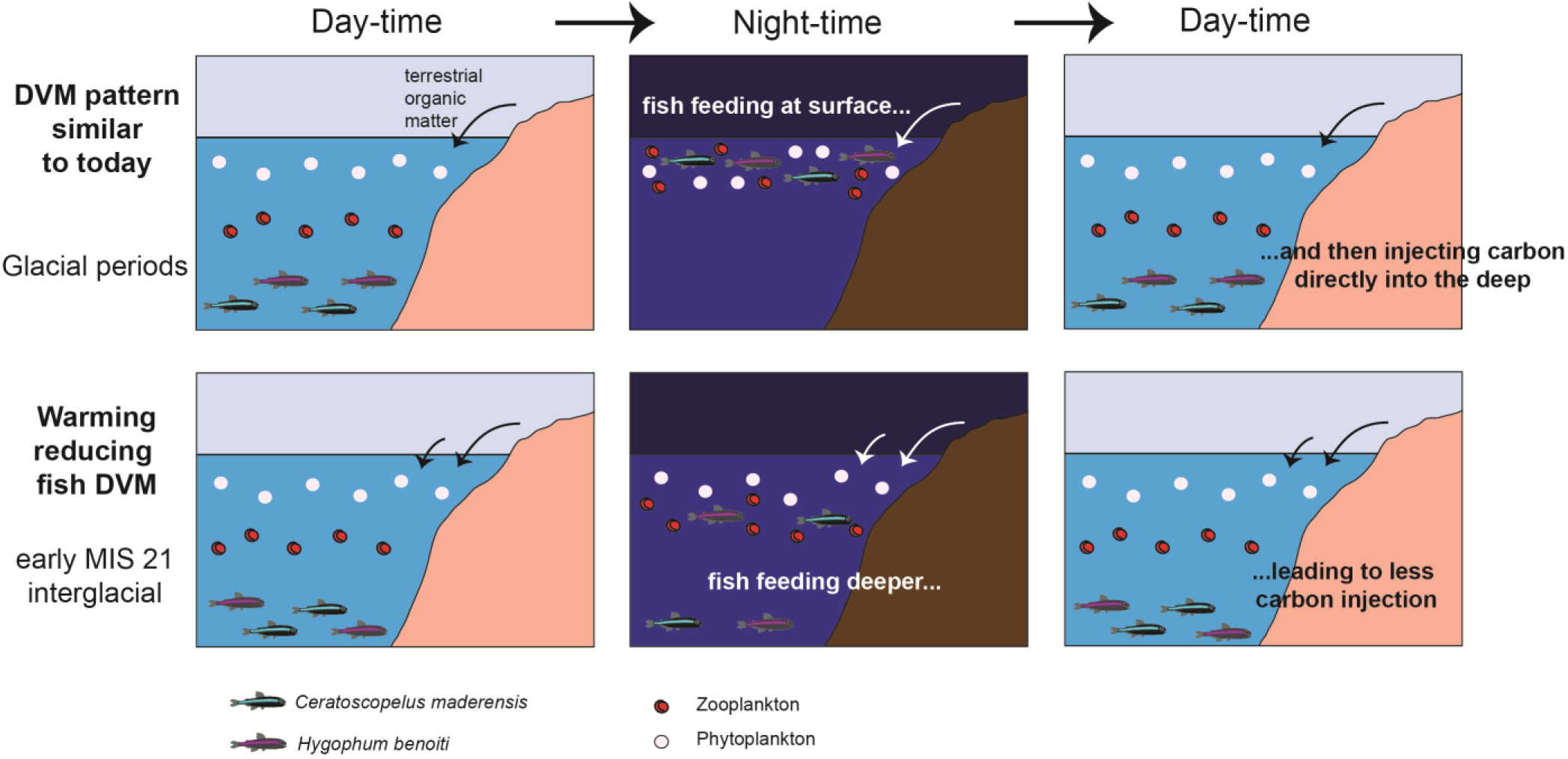
Schematic representation of the changes in diel vertical migration patterns during the study interval, as suggested by our data.

A drastic shift to greater lifetime-average depth (model estimate: 1502 m) is observed for *Ceratoscopelus maderensis* in early MIS 21 (Fig. 6). This suggest a slowing down of DVM and/or drastic shift of these fish to much greater mesopelagic depths than before (Fig. 8). These two alternative explanations would have opposite effects on the biological carbon pump and therefore on carbon sequestration and burial. In the central Aegean, Pallacks et al. (2025) evidenced a likely absence of myctophids at 10–7 ka BP, which they attributed to deoxygenation occurring during sapropel formation (core recovered at 559.2 m water depth). No sapropel was found in the Lardos sediments, which deposited at shallower paleodepths (Figs. 5 & 6). Moreover, there is no decline in absolute or relative abundance of myctophid otoliths in the sediments corresponding to interglacial periods of the Lindos Bay Formation (Agiadi et al., 2018, 2023). If myctophids expanded their depth distribution to greater depths, but continued to perform DVM as before, then we would expect to see an increase in organic carbon burial at the time, which is not evidenced by TOC or foraminiferal *δ*^13^C data (Fig. 3). In contrast, we support that the DVM capacity of fishes was hindered by the warming that occurred in the early part of MIS 21, resulting in a reduction of the biological carbon pump efficiency (Fig. 8) and relatively stable carbon burial despite the enhanced terrestrial input (increased BIT; Fig. 3) and subsequent interglacial. The rather great lifetime-average depths for the juveniles of both *C. maderensis* (457 m) and *H. benoiti* (933 m) around the middle part of MIS 21 reinforce the interpretation of a reduction in DVM, and further signal a gradual re-establishment of regular DVM patterns (replicate data confirming this are available in the dataset Agiadi et al. (2024b). This event was concurrent with a shift in global and Mediterranean SST (Fig. 7). Although higher resolution and more sites are necessary to investigate the extent of this phenomenon, our results support the hypothesis that warming reduces the biological carbon pump efficiency by restricting mesopelagic zone ecosystem functioning (Boscolo-Galazzo et al., 2021).

Teleost fishes are the largest vertebrate group, occupying all aquatic ecosystems, from high-altitude lakes to the abyssal plain of the oceans (FishBase, 2024; Nelson et al., 2016). They all have otoliths, which are nicely preserved in Mesozoic and Cenozoic sediments from polar to tropical latitudes (Martin and Dunn, 2000; Schwarzhans, 2018; Schwarzhans et al., 2017): even though aragonitic, otoliths enter the sediment protected, either within the decaying carcass of the fish or inside the fecal pellets of larger predators (Agiadi et al., 2022). Because their oxygen isotopic composition is correlated with those of foraminifera occupying the same depths, otolith *δ*^18^Ο is a promising paleoceanographic proxy (Agiadi et al., 2024a; Price et al., 2009). Establishing this proxy can facilitate global correlations, even in the absence of foraminifera, and including deep oceanic facies.

In order to use otolith oxygen isotopes as paleoceanographic proxies, some limitations must be considered: First, otoliths are no substitute for other proxies, in the sense that they can never provide the high resolution achieved through e.g. foraminifera. Second, it is of the utmost importance that the otoliths analyzed have been carefully identified, considering the great number of teleost species and the variety of environments they can occupy. Analyzing together otolith specimens of different or even unknown species creates problems and errors in interpretation, as their values can correspond to vastly different ocean depths (e.g. (Ivany et al., 2000). It is fundamental that the ecology of the species whose otoliths are analysed is well known, which is common-case for many species found in Late Cenozoic marine sediments, and many Early Cenozoic genera.

## Conclusions

We employed a multi-proxy approach to reconstruct the response of mesopelagic fish DVM to paleoceanographic changes that took place between 923 and 756 ka in the Eastern Mediterranean. Glacial–interglacial changes in the Pleistocene tested the limits of marine ecosystem resilience in multiple ways, resulting in distribution range shifts, ecophenotypic adaptations, and ultimately species extinctions (Agiadi et al., 2011, 2018, 2023; Porz et al., 2024). During the studied time interval, the Eastern Mediterranean was a warm subtropical area with fluctuating influxes of terrestrial organic matter. The global warm MIS 21 initiation, the sea-level drop in MIS 20, and high productivity accompanied by enhanced organic matter burial during MIS 19 are the most notable environmental changes we evidence in the study area. Across the food web, marine organisms seem to have been affected by the warming that took place during the early part of MIS 21, with increasing observed in the biomasses of the invertebrate groups (foraminifera, ostracods and sponges) and a reduction in DVM for mesopelagic fishes. Similarly, large variations are observed in foraminifera, ostracod and sponge biomasses within the MIS 19 interglacial. However, in contrast to MIS 21, we could not find evidence of a disruption in the fish DVM pattern in MIS 19, even though the fish abundances and body sizes have been shown to be affected by this interglacial (Agiadi et al., 2023). Although our ecosystem-level approach is challenging, it offers a holistic approach for assessing the impacts of paleoenvironmental changes that can be most informative for ecosystem management.

## Acknowledgements

This research was funded by the Austrian Science Fund (FWF) Grant DOI 10.55776/M2894 (P.I.: KA). For the purpose of open access, the author has applied a CC BY public copyright license to any Author Accepted Manuscript version arising from this submission. FQ was supported by the National programs Tellus-INTERRVIE of CNRS-INSU. AFA and JMT received funding from the Spanish project ProOceans (Ministerio de Ciencia e Innovación, Proyectos de I + D + I (RETOSPID2020-118097RB-I00)). The authors acknowledge institutional support of the ‘Severo Ochoa Center of Excellence’ accreditation (CEX2019-000928-S) and the Catalan government through the iMARES research group of quality at ICM-CSIC. We warmly thank Arnauld Vinçon-Laugier for technical assistance with the mass spectrometer, Uli Treffert for assistance in the SBiK-F laboratories, Jean-Jacques Cornée and Pierre Moissette for showing us the Lardos section, and George Kontakiotis and Vasiliki Lianou for their help in the field.

## Authors Contribution

Conceptualization: KA, FQ; Formal analysis: KA, IV, AV, AFA, JMT, FQ; Funding acquisition: KA, FQ; Investigation: KA, IV, AV, EK, SZ, AFA, JMT, FQ; Methodology: KA, IV, SZ, AFA, JMT, FQ; Supervision: KA, FQ; Writing – original draft: KA, IV, AV, SZ, AFA, JMT, FQ; Writing – review and editing: EK.

## Data and Code Availability Statement

The data produced for this work is publicly available as the dataset Agiadi et al. (2024b). The code to estimate the fish lifetime-average depth is publicly available in (Fuster-Alonso et al., 2024). A preprint version related to this manuscript is available at (Agiadi et al., 2024c)

