## Supplementary material for "Mesopelagic fish responses to Pleistocene climatic variability in the Eastern Mediterranean and implications for the biological pump"

### *Climate, productivity and organic carbon burial*

Four paleoceanographic regimes were identified within the studied time interval in the southeastern Aegean, which are bound by regime shifts, each shift related to a different factor (SST, depth, primary or secondary production), as explained below: a) 923–858 ka; b) 851–798 ka; c) 788–762 ka; and d) 760–756 ka.

*923–858 ka*

This interval starts from the earliest time represented by our studied sediments and last until the paleoceanographic regime shift recorded by $\delta^{13}C_{ruber}$ at 851 ka, which is interpreted here to correspond to a significant increase in primary productivity by phytoplankton afterwards. After 864 ka, the regime shift in the sponge weighted average size is coupled with a large increase in BIT (Fig.3 of main text) and foraminiferal biomasses, and a decrease in $\delta^{18}O_{inflata}$ and $\delta^{18}O_{peregrina}$ (while $\delta^{18}O_{ruber}$ remains at the same levels), suggesting lengthening of the thermal gradient in the water column associated with the influx of terrestrial organic matter (i.e., higher BIT) that benefited secondary production. The interval between 923 and 858 ka includes the MIS 23 interglacial, the MIS 22 glacial, and the early part of the MIS 21 interglacial. SST declines from MIS 23 to MIS 22 and increases in MIS 21 (Fig. 2). TOC (and TN) increase throughout the studied interval (Fig. 3). Planktonic and benthic foraminifera biomasses are low in MIS 23 and even lower in MIS 22, but increase drastically in MIS 21 (Fig. 4). Overall, stable high $\delta^{13}C_{bulk}$ values appear for this interval, and $\delta^{13}C_{peregrina}$ shows intermediate values (Fig. 3). $\delta^{13}C_{inflata}$ decreases in the 864–858 ka interval. In contrast, $\delta^{13}C_{ruber}$ values are higher in MIS 21, and lower in MIS 22 and MIS 23. Our results suggest that, at Lardos, productivity during the 923–864 ka interval fluctuates with climate, as reflected by the $\delta^{13}C_{ruber}$ values (Fig. 3) and the foraminiferal biomasses (Fig. 4). This is because TOC and $\delta^{13}C$ values of the bulk sediments and foraminiferal shells generally reflect changes in productivity and/or organic carbon burial (Jing et al., 2024; Spezzaferri, 1995). Particularly, the $\delta^{13}C$ of foraminifera is a proxy of the $\delta^{13}C$ of dissolved inorganic carbon in the seawater where shell calcification took place, which increases with primary productivity, influx of terrestrial organic matter and water mixing (Ravelo and Hillaire-Marcel, 2007). Additionally, benthic foraminiferal accumulation rates from deep-sea drilling sites are often used to estimate changes in productivity (Diester-Haass et al., 2018). In contrast, organic carbon burial declines during the 923–864 ka interval, since the benthic $\delta^{13}C_{peregrina}$ values remain approximately the same.

*851–798 ka*

The regime shift in $\delta^{13}C_{ruber}$ denotes a paleoenvironmental or ecosystem change at Lardos at 851 ka. Essentially, a drastic increase in primary productivity characterizes this interval, which is immediately reflected in planktonic and benthic foraminiferal biomasses, ostracod and sponge spicule counts, which are generally high in this interval (Fig. 4). However, the $\delta^{13}C_{inflata}$ and $\delta^{13}C_{peregrina}$ records do not pick up this shift, implying that surface waters were the most affected by the change than deeper waters were. Within the 851–798 ka interval, BIT increases to a maximum around the MIS 21/MIS 20 termination (Fig. 3), reflecting more terrestrial organic matter, and both *pfb* and *bfb* increase at the same time. This interval encompasses most of the MIS 21 interglacial, after the SST maximum at 858 ka (Fig. 2), and the MIS 20 glacial. Upwards, another important regime shift is detected at 788 ka in both $\delta^{13}C_{peregrina}$ (Fig. 2) and the ostracod weighted-average body size, and it is attributed to a decrease in secondary, benthic production after this time.

*788–762 ka*

The interval between 788 and 762 ka is characterized at Lardos by a regime shift in $\delta^{13}C_{peregrina}$ and the ostracod weighted-average body size, which are interpreted here to reflect a drop in secondary, benthic production. The $\delta^{13}C_{peregrina}$ (and the $\delta^{13}C_{bulk}$) values during this interval are generally the lowest, while ostracods show a drastic decline in body size with an almost complete absence of sizes > 500 µm. From 780 ka toward MIS 18, SST indicates a cooling trend (Fig. 2), which has been also detected along the Iberian margin (Atlantic) and has been there related to a progressive southward migration of the polar front (Girone et al., 2023). A decreasing trend is also evident in SSS (Fig. 2) from 780 to 762 ka. In contrast to ostracod body sizes, higher planktonic foraminifera biomasses are observed throughout the 788–762 ka interval. $\delta^{13}C_{ruber}$ drops in MIS 19 (Fig. 2). Therefore, we propose that higher surface, primary productivity during MIS 19 was accompanied by enhanced organic carbon burial and lower secondary, benthic production to explain the concurrent higher planktonic foraminifera biomass and low $\delta^{13}C_{peregrina}$ and $\delta^{13}C_{bulk}$ values. Higher primary productivity and organic carbon burial have been reported at least for the early part of MIS 19 in the Western Mediterranean as well (ODP Site 975; (Marino et al., 2022; Quivelli et al., 2020).

*760–756 ka*

In our study area, the paleoceanographic conditions change at the MIS 19/MIS 18 termination, as captured by a regime shift in TOC, $\delta^{18}O_{peregrina}$ and $\delta^{18}O_{ruber}$ at 760 ka. TOC shows low values (Fig. 3), whereas $\delta^{18}O_{peregrina}$ and $\delta^{18}O_{ruber}$ show higher values than in MIS 19 (Fig. 2). SST and SSS ranges are similar to MIS 19, but the estimated paleodepth is shallower at this time (Fig. 2). In SW Europe, winter precipitation and the East Asia summer monsoon strength peaked during MIS 18, contributing to the expansion of the Northern Hemisphere ice sheets (Sánchez Goñi et al., 2023). However, at Lardos, the shallowing of the area most likely explains the observed regime shift at 760 ka.

### ***Supplementary Figures and Tables***

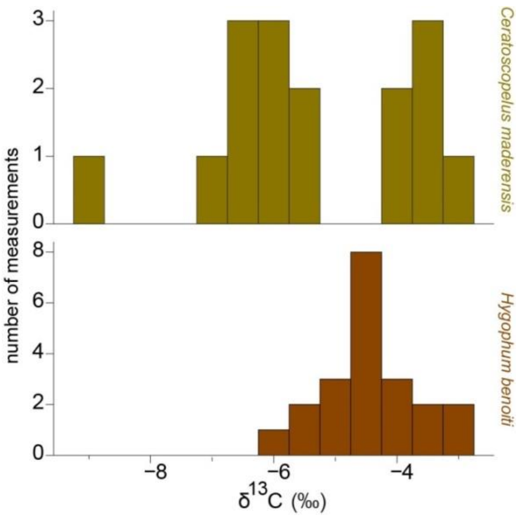


**Figure S1. Distribution of the $\delta^{13}C_{mader}$ and $\delta^{13}C_{benoiti}$ values.**

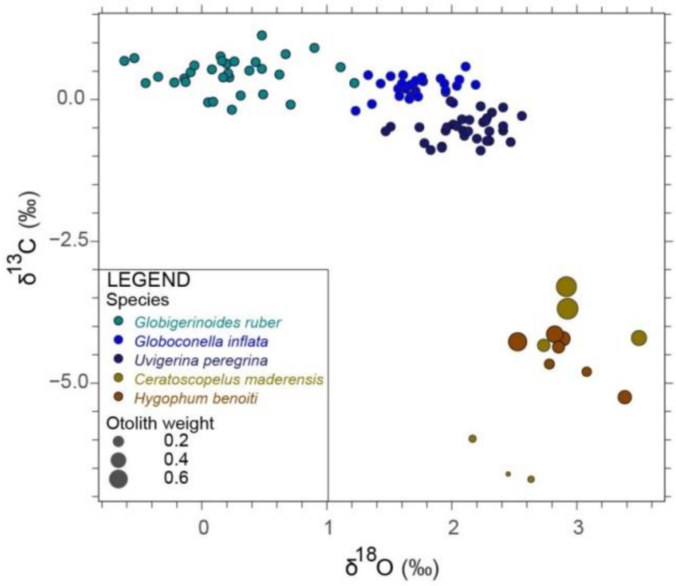


**Figure S2.** $\delta^{13}$**C vs.** $\delta^{18}$**O in foraminifera and the fish otoliths. The different species are clustered well, and otolith are clearly separated from foraminifera values. The circle diameter for the otoliths is scaled according to the otolith weight, which is used here as an indication of fish size, since otoliths grow throughout the lifetime of the fish. Therefore, within the same species, small (and light) otoliths correspond to younger individuals than larger (and heavier) ones.**

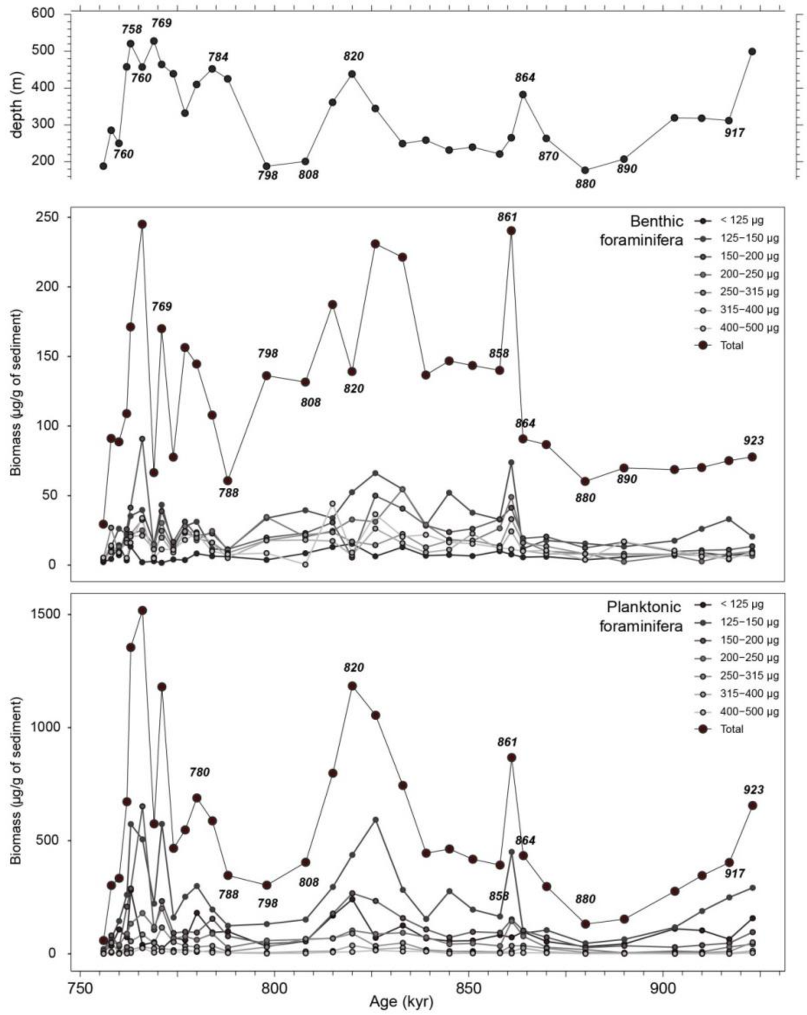


**Figure S3. Contribution of the different size classes to total benthic and planktonic foraminifera biomass.**

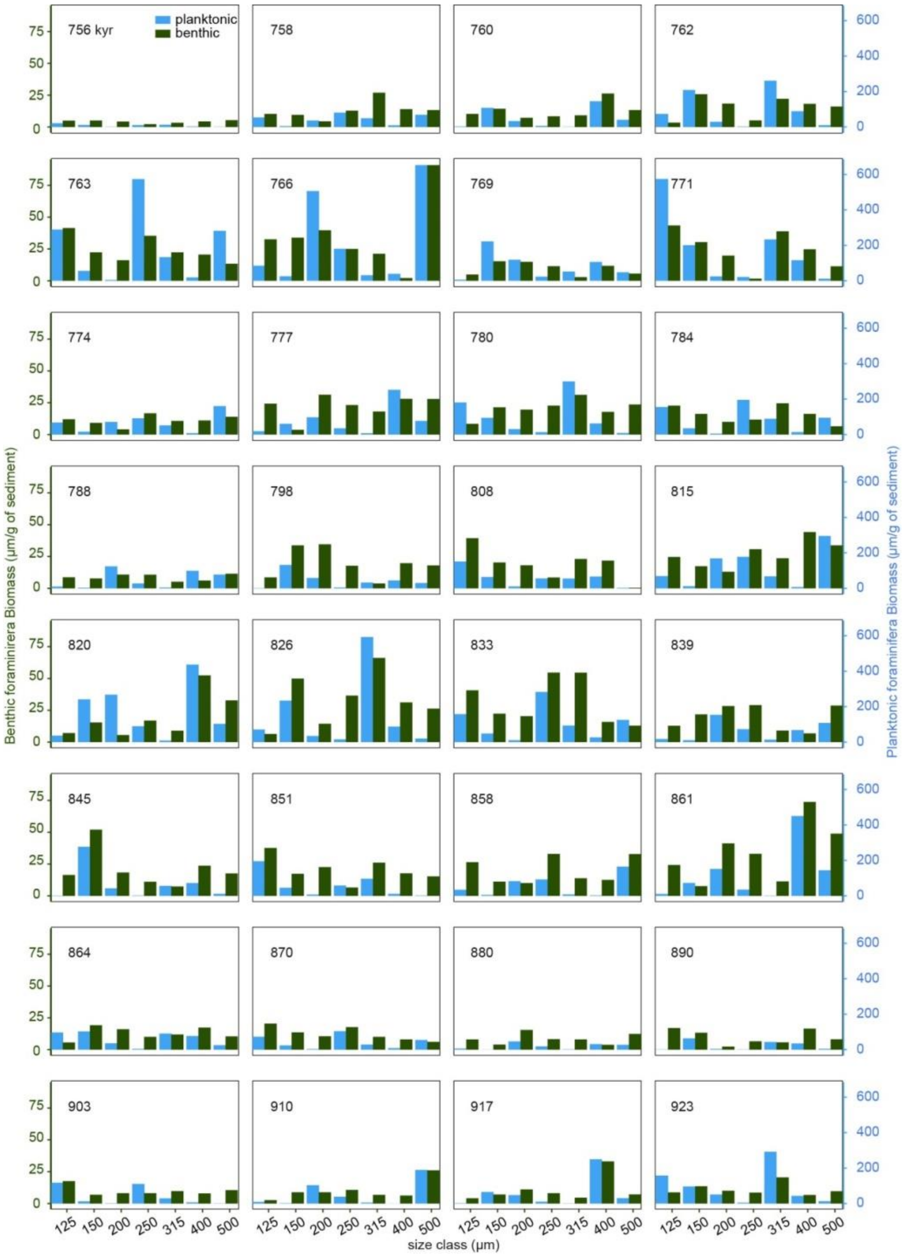


**Figure S4. Benthic and planktonic foraminifera biomass at Lardos by size class through time.**

**Table S1. Mean SST values estimated at different in the region for each MIS.**

| MIS | Site | SST (°C) | Proxy | Reference |
|---|---|---|---|---|
| 18 | Lardos | 23.5 | $TEX_{86}$ | This study |
| | M.J. | 15.0 | $U^{k'}_{37}$ | (Marino et al., 2020) |
| | LC07 | 20.7 | $U^{k'}_{37}$ | (Martínez-Dios et al., 2021) |
| | U1385 | 15.6 | $U^{k'}_{37}$ | (Rodrigues et al., 2017) |
| 19 | Lardos | 22.8 | $TEX_{86}$ | This study |
| | LC07 | 19.8 | $U^{k'}_{37}$ | (Martínez-Dios et al., 2021) |
| | ODP975 | 16.3 | Forams | (Quivelli et al., 2020) |
| | U1385 | 18.1 | $U^{k'}_{37}$ | (Rodrigues et al., 2017) |
| 20 | Lardos | 23.7 | $TEX_{86}$ | This study |
| | LC07 | 18.9 | $U^{k'}_{37}$ | (Martínez-Dios et al., 2021) |
| | ODP975 | 10.8 | $U^{k'}_{37}$ | (Quivelli et al., 2021) |
| | U1385 | 14.0 | $U^{k'}_{37}$ | (Rodrigues et al., 2017) |
| 21 | Lardos | 22.2 | $TEX_{86}$ | This study |
| | LC07 | 19.1 | $U^{k'}_{37}$ | (Martínez-Dios et al., 2021) |
| | U1385 | 16.9 | $U^{k'}_{37}$ | (Rodrigues et al., 2017) |
| 22 | Lardos | 23.6 | $TEX_{86}$ | This study |
| | LC07 | 16.9 | $U^{k'}_{37}$ | (Martínez-Dios et al., 2021) |
| | U1385 | 11.4 | $U^{k'}_{37}$ | (Rodrigues et al., 2017) |
| 23 | Lardos | 23.5 | $TEX_{86}$ | This study |
| | LC07 | 18.7 | $U^{k'}_{37}$ | (Martínez-Dios et al., 2021) |
| | U1385 | 15.5 | $U^{k'}_{37}$ | (Rodrigues et al., 2017) |

#### *Supplementary References*


Diester-Haass, L., Billups, K., and Lear, C.: Productivity changes across the mid-Pleistocene climate transition, Earth-Sci. Rev., 179, 372–391, https://doi.org/10.1016/j.earscirev.2018.02.016, 2018.

Girone, A., De Astis, A., Sierro, F. J., Hernández-Almeida, I., Garcia, M. A., Sánchez Goñi, M. F., Maiorano, P., Marino, M., Trotta, S., and Hodell, D.: Planktonic foraminifera response to orbital and millennial-scale climate variability at the southern Iberian Margin (IODP Site U1385) during Marine Isotope Stages 20 and 19, Palaeogeogr. Palaeoclimatol. Palaeoecol., 615, 111450, https://doi.org/10.1016/j.palaeo.2023.111450, 2023.

Jing, Y., Zhang, T., Zhu, B., Zhao, J., Zhao, X., Dou, Y., Li, Q., Cai, F., Hu, B., and Dong, L.: Organic carbon burial and their implication on sea surface primary productivity in the middle Okinawa Trough over the past 200 ka, Front. Mar. Sci., 11, https://doi.org/10.3389/fmars.2024.1331940, 2024.

Marino, M., Girone, A., Gallicchio, S., Herbert, T., Addante, M., Bazzicalupo, P., Quivelli, O., Bassinot, F., Bertini, A., Nomade, S., Ciaranfi, N., and Maiorano, P.: Climate variability during MIS 20–18 as recorded by alkenone-SST and calcareous plankton in the Ionian Basin (central Mediterranean), Palaeogeogr. Palaeoclimatol. Palaeoecol., 560, 110027, https://doi.org/10.1016/j.palaeo.2020.110027, 2020.

Marino, M., Rodrigues, T., Quivelli, O., Girone, A., Maiorano, P., and Bassinot, F.: Paleoproductivity proxies and alkenone precursors in the Western Mediterranean during the Early-Middle Pleistocene transition, Palaeogeogr. Palaeoclimatol. Palaeoecol., 601, 111104, https://doi.org/10.1016/j.palaeo.2022.111104, 2022.

Martínez-Dios, A., Pelejero, C., Cobacho, S., Movilla, J., Dinarès-Turell, J., and Calvo, E.: A 1-Million-Year Record of Environmental Change in the Central Mediterranean Sea From Organic Molecular Proxies, Paleoceanogr. Paleoclimatology, 36, e2021PA004289, https://doi.org/10.1029/2021PA004289, 2021.

Quivelli, O., Marino, M., Rodrigues, T., Girone, A., Maiorano, P., Abrantes, F., Salgueiro, E., and Bassinot, F.: Surface and deep water variability in the Western Mediterranean (ODP Site 975) during insolation cycle 74: High-resolution calcareous plankton and molecular biomarker signals, Palaeogeogr. Palaeoclimatol. Palaeoecol., 542, 109583, https://doi.org/10.1016/j.palaeo.2019.109583, 2020.

Quivelli, O., Marino, M., Rodrigues, T., Girone, A., Maiorano, P., Bertini, A., Niccolini, G., Trotta, S., and Bassinot, F.: Multiproxy record of suborbital-scale climate changes in the Algero-Balearic Basin during late MIS 20 - Termination IX, Quat. Sci. Rev., 260, 106916, https://doi.org/10.1016/j.quascirev.2021.106916, 2021.

Ravelo, A. C. and Hillaire-Marcel, C.: The use of oxygen and carbon isotopes of foraminifera in Paleoceanography, in: C. Hillaire-Marcel, A. de Vernal, Proxies in Late Cenozoic Paleoceanography, Elsevier, Amsterdam, 735–764, 2007.

Rodrigues, T., Alonso-García, M., Hodell, D. A., Rufino, M., Naughton, F., Grimalt, J. O., Voelker, A. H. L., and Abrantes, F.: A 1-Ma record of sea surface temperature and extreme cooling events in the North Atlantic: A perspective from the Iberian Margin, Quat. Sci. Rev., 172, 118–130, https://doi.org/10.1016/j.quascirev.2017.07.004, 2017.

Sánchez Goñi, M. F., Extier, T., Polanco-Martínez, J. M., Zorzi, C., Rodrigues, T., and Bahr, A.: Moist and warm conditions in Eurasia during the last glacial of the Middle Pleistocene Transition, Nat. Commun., 14, 2700, https://doi.org/10.1038/s41467-023-38337-4, 2023.

Spezzaferri, S.: Planktonic foraminiferal paleoclimatic implications across the Oligocene-Miocene transition in the oceanic record (Atlantic, Indian and South Pacific), Palaeogeogr. Palaeoclimatol. Palaeoecol., 114, 43–74, https://doi.org/10.1016/0031-0182(95)00076-X, 1995.
